# Neural encoding of grasp and object properties in the posterior parietal and motor cortices of the cortical grasping network in tetraplegic humans

**DOI:** 10.64898/2026.09.10.750758

**Authors:** Mackenzie J. Thurston, David Bjånes, Sarah Wandelt, Kelsie Pejsa, Brian Lee, Charles Liu, Richard A. Andersen

## Abstract

The cortical grasping network (CGN) is responsible for the ability to physically interact with the world around us using our hands. For patients with motor impairments due to neurogenerative disease or traumatic injury, brain-machine interfaces (BMIs) offer a potential pathway towards restoration of dexterous hand control via recording neural activity throughout the CGN. However, grasping objects with robotic BMI devices has proved challenging, when utilizing neural signals from only motor cortex (MC, a subregion of the CGN). It is currently unclear how the presence of an object might compromise BMI performance; thus this work explores the interactive neural representation of grasp and objects throughout the CGN.

Three tetraplegic human participants performed grasp motor imagery during imagined object manipulation while we recorded neural activity from the supramarginal gyrus (SMG), anterior intraparietal cortex (AIP), motor cortex (MC), and primary somatosensory cortex (S1). All regions within the CGN represented whole hand configuration of imagined grasps during motor planning and imaged execution. Additionally, grasp-related neural activity in each region was modulated by context (motor planning vs. imagined execution and object present vs. not present). SMG and AIP represented object shape during motor planning. PPC encoded both grasp and object properties simultaneously from mostly unique subpopulations of neurons. This separability in higher cortical regions could be a critical for stable BMI grasp performance.

## INTRODUCTION

Most activities of daily living (ADLs), a commonly used clinical indicator of a person’s ability to care for themselves independently, require the ability to grasp^1^. For people with tetraplegia, these activities become incredibly difficult, if not entirely impossible. In a survey of people with Spinal Cord Injuries (SCI), when asked what would provide the largest improvement in quality of life, people with tetraplegia identified regaining hand and arm function as their top priority^2^.

Brain-Machine Interfaces (BMIs) offer a unique opportunity to restore motor function to SCI individuals. BMIs operate by circumventing the injury in the spinal cord and harnessing information directly from neural activity in healthy, functional cortex to control robotic limbs and restore hand movement^3^. Intracortical BMIs use microelectrodes implanted directly into the cortex to record spiking activity at the level of single neurons with high signal-to-noise ratio (SNR) and high spatial and temporal resolution^4^. While there are many forms of BMIs, these qualities make intracortical BMIs an ideal candidate for neuroprosthetics.

The cortical grasping network (CGN), the brain regions responsible for the ability to grasp, are the top candidates for microelectrode implantation sites for a grasp BMI. The CGN includes regions of the posterior parietal cortex (PPC), such as the supramarginal gyrus (SMG) and anterior intraparietal cortex (AIP), and motor cortex (MC). From non-human primate (NHP) research^5–8^, MC is a primary motor area encompassing grasp and reach regions^9,10^. Yan et al. (2026) recently demonstrated MC may contain object-specific information during grasping^11^. PPC is known to be an integration site for sensorimotor transformations and movement intention^12^. Along with encoding grasp and reach^8,13^, AIP may also contribute to visual processing of object sizes and shapes^5^.

Although many organizational principles of grasp and reach representation may be conserved between NHP and human MC^14,15^ and AIP^8,16^, human CGN exhibits important specializations, including the expansion of posterior parietal cortex and the emergence of cortical regions without clear NHP homologs, such as SMG. Imaging and electrophysiologic studies have implicated SMG in tool use^17–19^, grasping and manipulating objects^20,21^, reaching^22^, and semantic representation^23^. With such heterogeneous neural representations, this rich mixture of variables may be uniquely advantageous to utilize for a grasp BMI device.

Motor cortex has widely investigated for motor BMI applications^3,24,25^, with a recent demonstration of reach and hand aperture control (open or closed) enabled a participant to complete a box and blocks task^26^. However, restoring dexterous hand function to support a wide range of hand shapes and grasp configurations will require understanding how object characteristics are represented throughout the cortex, as object characteristics are key determinants of grasp selection. Prior studies found the presence of an object in the visual field drastically decreased the performance of their BMI such that participants could no longer consistently grasp using a robotic limb^27^. While computational methods could mitigate such performance loss, overall accuracy was significantly decreased relative to the initial decoding performance without the object present.

This work investigated neural representation of grasp and object throughout human CGN by recording single neuron activity from three chronically implanted participants during a grasp imagery task. We dissociated object characteristics (size and shape) from intended grasp to investigate how object presence altered neural representation. These findings have the potential to inform the development of a functional grasp BMI.

## RESULTS

We found differential neural representations across the human cortical grasping network (CGN) for motor planning and imagined execution, as well as object property variables during imagined grasping of objects. Across three human participants (**Fig. 1A**), we recorded neural activity from thirteen chronically implanted microelectrode arrays, spanning supramarginal gyrus (SMG), post-central gyrus (S1), anterior intraparietal area (AIP), and the hand region of precentral gyrus in the motor cortex (MC). Both single neuron and population-level dynamics revealed distinct, complementary roles for posterior parietal, motor, and sensory cortices (**Fig. 2**).

**Figure 1.**
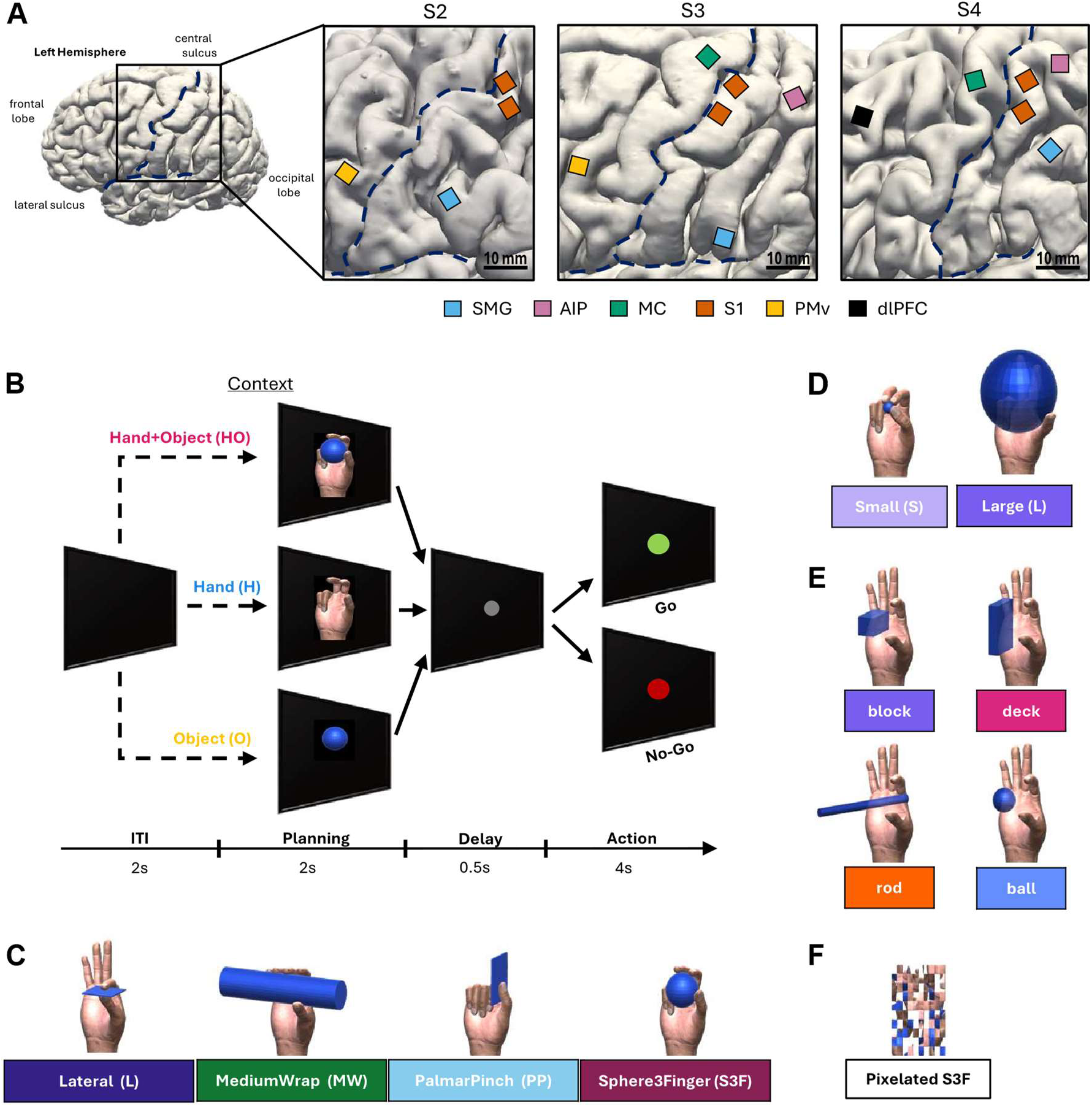
Experimental Design. A) Array locations for all three participants (S2, S3, S4). B) Trial Structure. Four phases: Inter-trial Interval (ITI), Planning, Delay, and Action. Three Cue Contexts: Hand+Object (HO), Hand only (H), and Object only (O). The color of the dot in Action represents Go (green) and No-Go (red) trials. C) Four grasp types used throughout all tasks. Grasp Types: Lateral (L), Medium Wrap (MW), Palmar Pinch (PP), and Sphere3Finger (S3F). D) Aperture size in the Varied Sizes Task: Small (S) and Large (L), S3F used as example. E) Disassociation Task: each grasp is paired with each object (for example, Lateral grasp shown with each object shape). Objects are modified to fit within all grasp types (see Methods). F) Pixelated Control Task: pixelated images of each grasp (for example, S3F grasp) used as a visual control.

**Figure 2.**
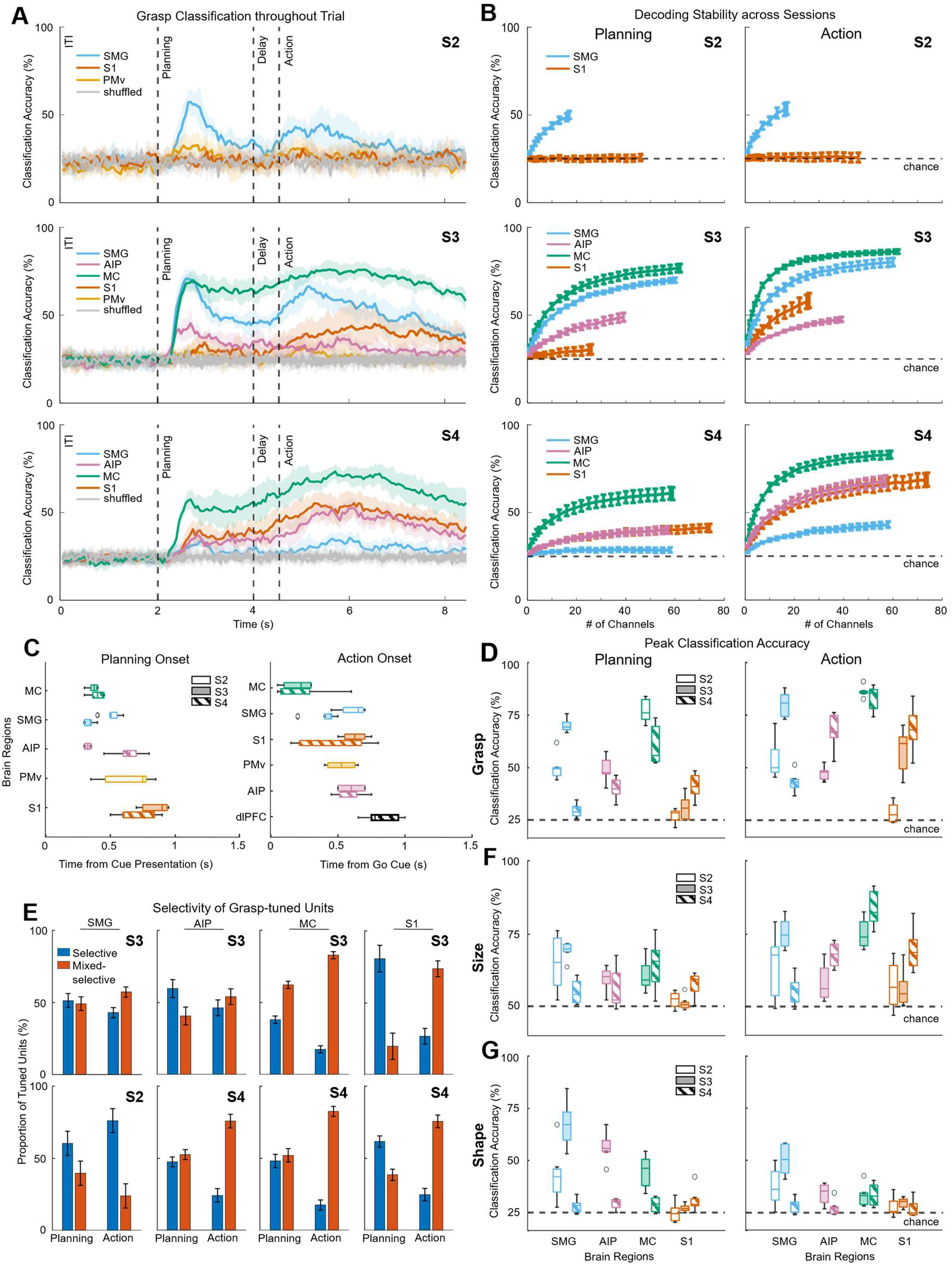
Grasp network. A) Session-averaged classification of grasp using an LDA classifier applied to each brain region within the cortical grasping network for 3 participants. B) Neuron Dropping Curves calculated using linear regression for each participant during Planning (column 1; S2: SMG: 50%, S1: 26%; S3: SMG: 70%, MC: 77%, AIP: 49%, S1: 30%; S4: SMG: 29%, MC: 61%, AIP: 40%, S1: 41%) and Action (column 2; S2: SMG: 54%, S1: 26%; S3: SMG: 80%, MC: 86%, AIP: 48%, S1: 58%; S4: SMG: 43%, MC: 83%, AIP: 68%, S1: 69%) phases. C) Planning latency (left) for each region determined by the onset of significant grasp decoding above chance during cue presentation. Planning latencies averaged across participants: MC: 383ms, SMG: 425ms, AIP: 483ms, PMv: 640ms, S1: 764ms. Execution latency (right) of each region determined by the onset of significant trial type (Go vs. No-Go trials) decoding above chance. Execution latencies averaged across participants: MC: 211ms, SMG: 511ms, S1: 520ms, PMv: 525ms, AIP: 589ms, dlPFC: 817ms. Regions are organized from shortest average latency (top) to longest average latency (bottom). See Methods for protocol details. D) Peak grasp classification accuracy of each brain region calculated from the Neuron Dropping Curves during Planning (left) and Action (right) phases. E) Selective vs. Mixed-selective grasp-tuned units during Planning and Action for each region. Example shown for Hand+Object, see Supplementary Fig. 5 for selectively during each context. F) Peak size classification accuracy of each brain region calculated from the Neuron Dropping Curves during Planning (left) and Action (right) phases. G) Peak shape classification accuracy of each brain region calculated from the Neuron Dropping Curves during Planning (left) and Action (right) phases.

To dissociate object properties from motor variables during planning and imagined execution, we utilized a cued, delayed motor task (**Fig. 1B**), instructing participants to imagine grasping objects. Sets of unique objects (ball, block, rod, card) and unique hand shapes (precision and power grasps) were selected from the “human grasping database”^28^ to cover the range of grasp taxonomies. With this paradigm, we manipulated the size, shape, and context of each object and motor action: 1) performing grasps with/without objects, 2) grasping the same object in a variety of sizes, and 3) grasping the same object with a variety of grasps (**Fig. 1C,D,E**). *We hypothesized motor and sensory regions would encode fundamental variables of motor control (grasp type, hand aperture) remaining invariant to object manipulation, while posterior regions would preferentially encode object properties (shape, size). Given supramarginal gyrus’s implication in tool use and grasping, we hypothesized SMG would contain both motor variables and object properties*.

Across the CGN, we found grasp type was strongly encoded in motor cortex and PPC. However, only specific regions of the CGN selectively encoded object size (**Fig. 2F**) and object shape (**Fig. 2G**). While SMG and AIP demonstrated robust shape encoding during motor planning independent of grasp type, MC largely encoded hand aperture (**Fig. 5P**). Object shape encoding persisted in SMG during imagined execution, albeit not as robustly (accuracy: 44%, chance: 25%) (**Fig. S2B**). Object size was immediately available in SMG during motor planning and continued to be decodable during imagined execution, along with MC (**Fig. 3P**, **Fig. 5P**). The recorded neural population within AIP did not appear to encode object size (**Fig. 4P**).

**Figure 3.**
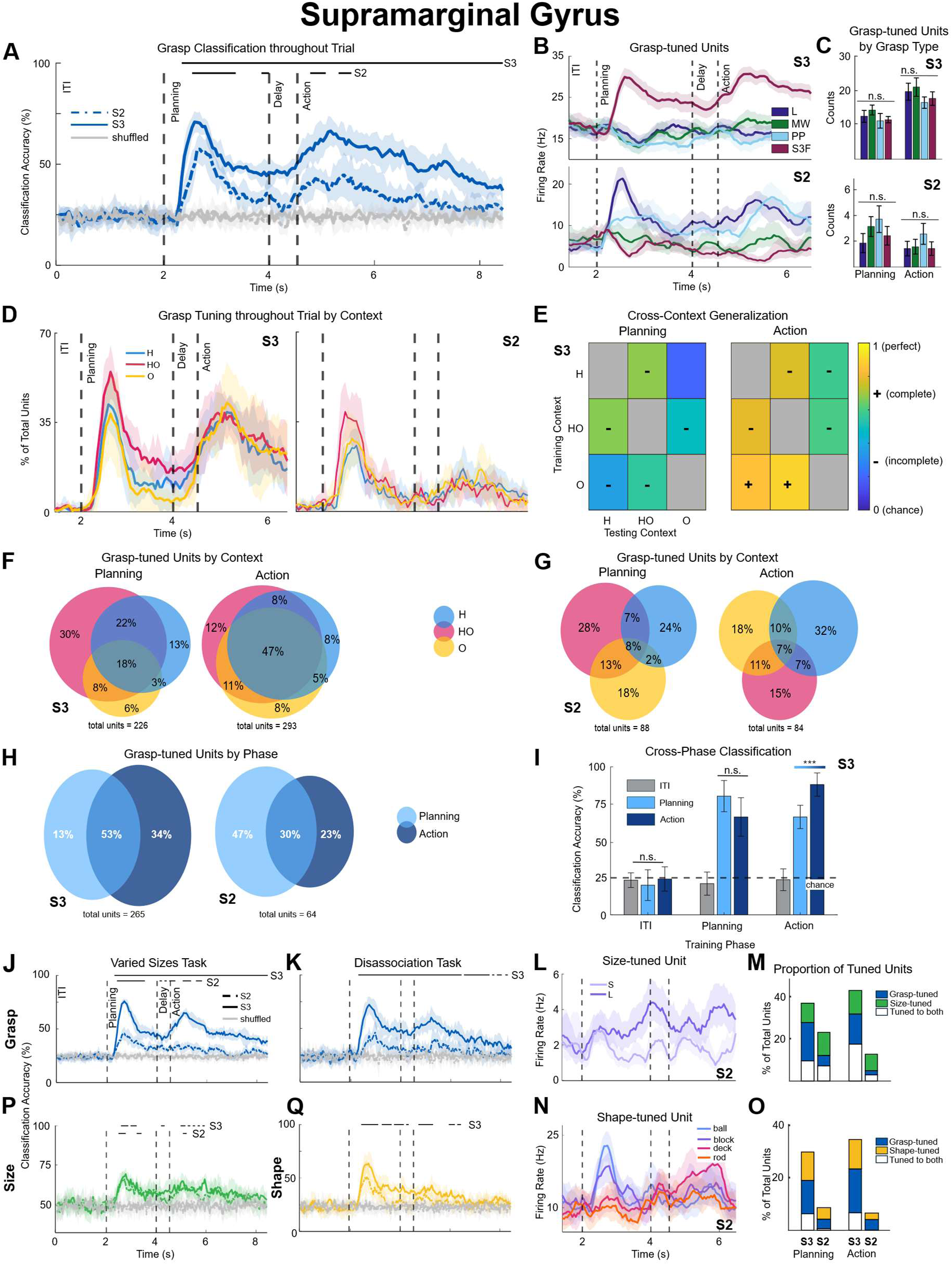
SMG exhibits partial mixed selectivity. A) Session-averaged classification of grasp using an LDA classifier is significantly above chance during motor planning and imagined execution across both participants. Classification performed at each 50ms timebin, mean accuracy (bold line) averaged across sessions with 95% standard error of the mean (SEM) confidence intervals. B) Grasp-tuned units in SMG. Top: an example unit tuned to one specific grasp type (S3F) during motor planning and imagined execution from participant S3. Bottom: an example unit tuned to two grasp types (L & PP) during planning and imagined execution from participant S2. C) Number of units tuned to specific grasp types averaged across sessions for each participant. No significant difference in number of units tuned to each grasp type. Data from Hand+Object (HO) context, see Supplementary Fig. 7 for all contexts. D) Percentage of units tuned to grasp across time within each context. Both participants demonstrate increased tuning when the hand and the object are shown simultaneously. E) Evaluating shared information across each context through cross-decoding analysis. Predominantly incomplete generalization among contexts during both Planning and Action. 0 = chance level generalization (dark blue), 1 = perfect generalization (yellow), calculated independently for each context/row. Gray = within context. - = incomplete generalization, + = complete generalization. Results from S3 only (see Methods). F/G) Proportion of grasp-tuned units in each context and the overlap among contexts. HO recruits the most units. H) Proportion of grasp-tuned units during Planning (light blue) and Action (dark blue) for two participants (S3, S2). There is substantial overlap between phases and the recruitment of many grasp-tuned units during Action only. HO results shown as example and results are largely consistent across contexts (see Supplementary Fig. 8). I) Evaluating shared information across each phase (Intertrial interval (ITI), Planning, Action). Cross-decoding generalized between Planning and Action phases, with complete generalization from Planning to Action and incomplete generalization from Action to Planning. Example shown for HO and is consistent across contexts (see Supplementary Fig. 9). Results from S3 only (see Methods). J/K) Session-averaged classification of grasp using an LDA classifier is significantly above chance during motor planning and imagined motor execution across both participants in both task variations. Grasp can still be accurately classified when manipulating object properties, as the accuracies and timing structure are nearly identical to the original task. L) A size-tuned example unit is shown where the unit’s firing rate is higher for an object that is Large and roughly baseline level response for Small throughout the trial. M) Proportion of units tuned to grasp (blue), size (green), or both (gray) for each participant during Planning (left) and Action (right) phases. A similar proportion of units are tuned to size during Planning and Action. Roughly half of the units are uniquely tuned to object size. N) An example unit is shown that is tuned to ball and block, responding only during motor planning. O) Proportion of units tuned to grasp (blue), shape (yellow), or both (gray) for each participant during Planning (left) and Action (right) phases. There is a unique shape population comparable to grasp-tuned units with little overlap. P) Size is significantly classified during Planning and Action in both participants at the timebin-level. Q) Shape is significantly classified during Planning in one participant and trending in another at the timebin-level.

**Figure 4.**
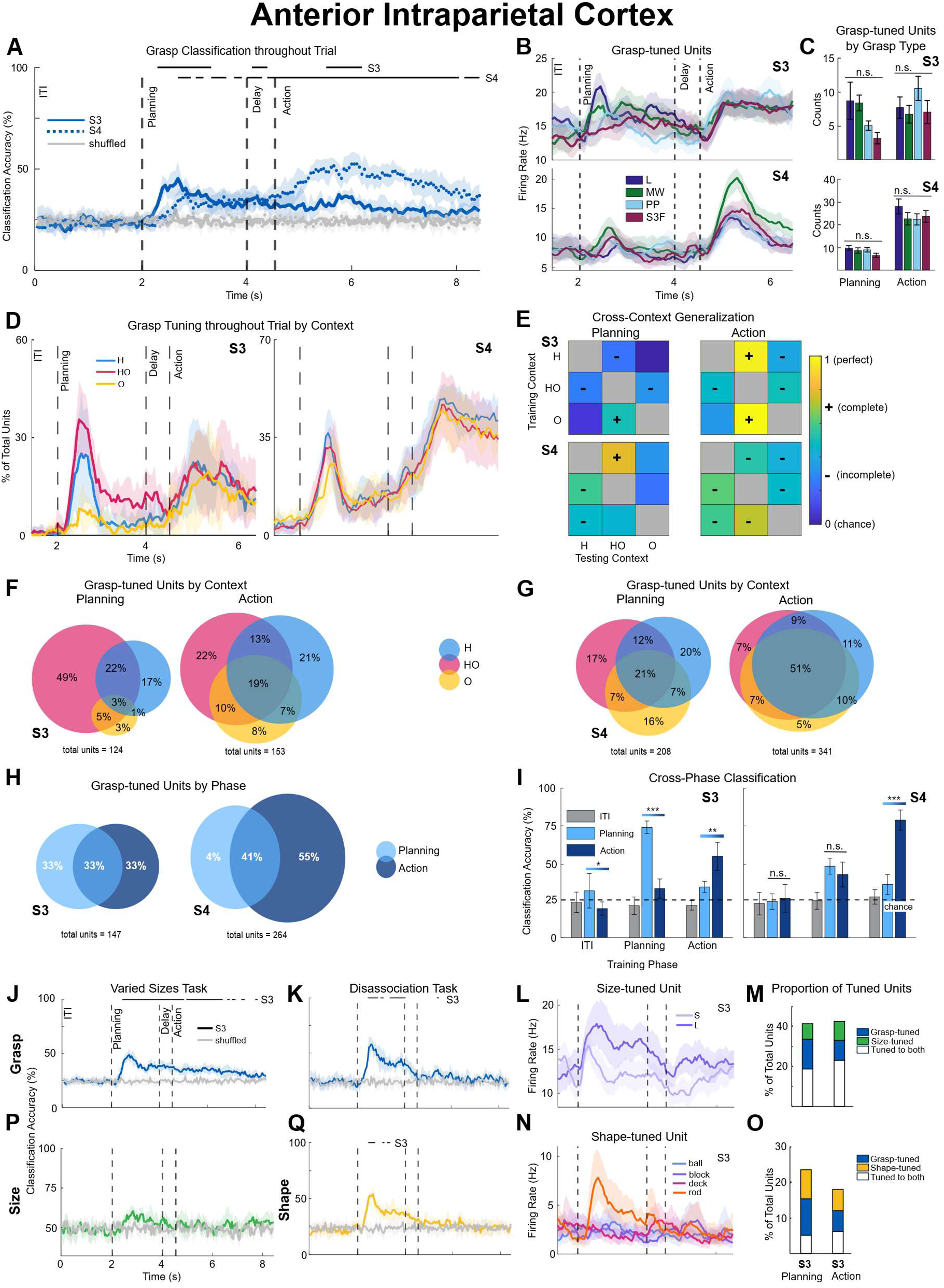
AIP demonstrates context-dependent encoding. A) Session-averaged classification of grasp using an LDA classifier is significantly above chance during motor planning and imagined execution across both participants. B) Grasp-tuned units in AIP. Top: an example unit demonstrating varying firing rates depending on the grasp type during motor planning, with a uniform response across all grasp types during imagined execution from participant S3. Bottom: an example unit with unique tuning to one grasp (MW) during Action from participant S4. C) Number of units tuned to specific grasp types averaged across sessions. No significant difference in number of units tuned to each grasp type. Data from Hand+Object (HO) context, see Supplementary Fig. 7 for all contexts. D) Percentage of units tuned to grasp across time within each context. Both participants demonstrate decreased tuning during Planning when the object is shown alone. Proportions during Action are similar across contexts. E) Evaluating shared information across each context. During Planning, context generalization was weaker than during Action. When generalization did occur, it was predominantly incomplete. 0 = chance level generalization (dark blue), 1 = perfect generalization (yellow), calculated independently for each context/row. Gray = within context. - = incomplete generalization, + = complete generalization. F/G) Evaluating the grasp-tuned units in AIP in each context and how these units overlap among contexts. HO recruits the most units. Overlap among all three contexts increases during Action. H) Number and proportion of units tuned to grasp during Planning (light blue) and Action (dark blue) for two participants (S3, S4). There is a large amount of overlap between phases and the recruitment of many grasp-tuned units during Action only. HO results shown as example and results are largely consistent across contexts (see Supplementary Fig. 8). I) Evaluating shared information across each phase (Intertrial interval (ITI), Planning, Action). Cross-phase classification shows incomplete generalization between Planning and Action phases. Example shown for HO and is consistent across contexts (see Supplementary Fig. 9). J/K) Session-averaged classification of grasp using an LDA classifier is significantly above chance during motor planning and imagined motor execution in both task variations. Grasp can still be accurately classified when manipulating object properties, though classification during Action largely drops out when object shape was manipulated. S4’s data was not included in these subsequent analyses (see Discussion). L) A size-tuned example unit is shown where the unit’s firing rate is higher for an object that is Large and lower for Small, but both invoke similar response patterns (increasing and decreasing at similar times). M) Proportion of units tuned to grasp (blue), size (green), or both (gray). A similar proportion of units are tuned to size during Planning and Action. Largely overlaps with grasp-tuned units. N) An example unit is shown that is selectively tuned to rod, responding only during motor planning. O) Proportion of units tuned to grasp (blue), shape (yellow), or both (gray). The shape-tuned population is comparable to grasp-tuned population with a mix of overlapping and uniquely shape-tuned units. P) Size is not significantly classified at any point in the trial at the timebin-level. Q) Shape is significantly classified during Planning at the timebin-level.

**Figure 5.**
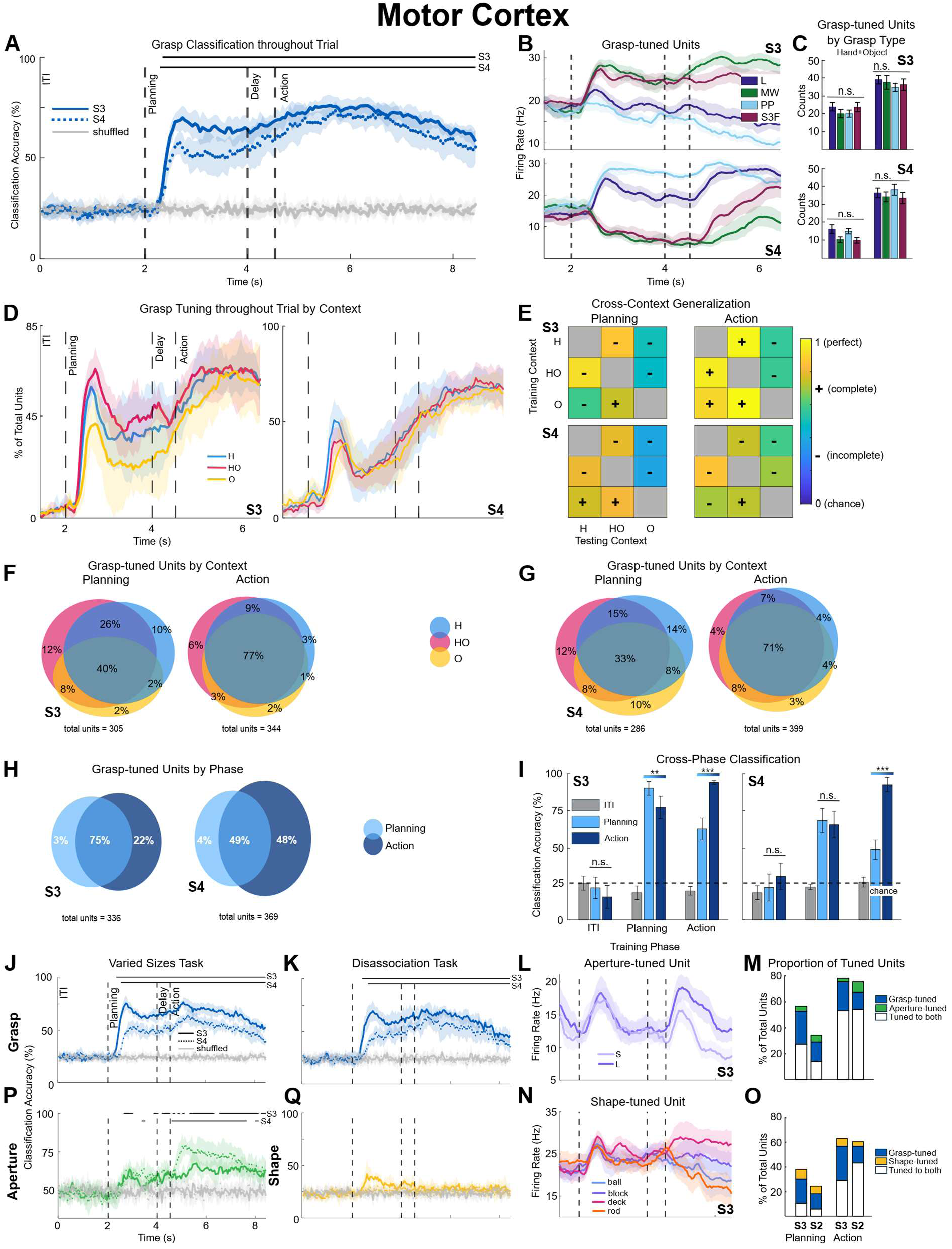
MC is highly selectively for motor-related encoding. A) Session-averaged classification of grasp using an LDA classifier is significantly above chance during motor planning and imagined execution across both participants. B) Grasp-tuned units in MC. Top: an example unit with selective tuning to two grasp types during motor planning, and unique tuning to each grasp type during imagined execution from participant S3. Bottom: a unit from participant S4 that demonstrates a similar grouping pattern of grasp types during Planning and unique firing rate patterns during Action. C) Number of units tuned to specific grasp types averaged across sessions. No significant difference in number of units tuned to each grasp type. Data from Hand+Object (HO) context, see Supplementary Fig. 7 for all contexts. D) Percentage of units tuned to grasp across time within each context. Both participants demonstrate similar proportions of tuning, with contexts involving a hand recruiting more units. E) Evaluating shared information across each context. During Planning and Action, contexts involving the hand (H & HO) reach near perfect generalization levels (yellow), with complete generalization during Action, compared to incomplete generalization to the object alone (O). 0 = chance level generalization (dark blue), 1 = perfect generalization (yellow), calculated independently for each context/row. Gray = within context. - = incomplete generalization, + = complete generalization. F/G) Evaluating the grasp-tuned units in each context and how these units overlap among contexts. Most units demonstrate extensive overlap, meaning they are tuned to grasp during all three contexts during both phases. H) Number and proportion of units tuned to grasp during Planning (light blue) and Action (dark blue) for two participants. There is a large amount of overlap between phases and the recruitment of many grasp-tuned units during Action only. HO results shown as example and results are largely consistent across contexts (see Supplementary Fig. 8). I) Evaluating shared information across each phase (Intertrial interval (ITI), Planning, Action). Planning and Action phases incompletely generalize to each other and demonstrate phase-dependent neuronal modulation. Example shown for HO and is consistent across contexts (see Supplementary Fig. 9). J/K) Session-averaged classification of grasp using an LDA classifier is significantly above chance during motor planning and imagined motor execution across both participants in both task variations. Grasp can still be accurately classified when manipulating object properties, as the accuracies and timing structure are nearly identical to the original task. L) An example unit demonstrates an equivalent response to both apertures early in motor planning but diverge during imagined execution with the Large aperture resulting in an increased firing rate relative to the Small. M) Proportion of units tuned to grasp (blue), hand aperture (green), or both (gray). A very small proportion of units are uniquely tuned to aperture compared to grasp, with a substantial increase from Planning to Action. Size-tuned units almost entirely overlap with grasp-tuned units. N) An example unit demonstrates varied firing rates depending on the object shape during imagined execution, with a near uniform response across all grasp types during motor planning. O) Proportion of units tuned to grasp (blue), shape (yellow), or both (gray). A smaller proportion of units are uniquely tuned to aperture compared to grasp, with a modest increase from Planning to Action. Most of the shape-tuned units overlap with grasp-tuned units. P) Aperture is significantly classified during Planning but predominantly during Action in both participants at the timebin-level. Q) Shape is not significantly classified during Planning or Action in either participant at the timebin-level.

### Grasp shape and object properties are decodable from all regions of the cortical grasping network

In all three participants, grasp type was broadly represented across the CGN, with MC consistently demonstrating the strongest encoding, followed closely by SMG, then AIP (**Fig. 2A,B**). Individual grasp types were equally well classified across areas (**Fig. S3**) and session to session representational stability was maintained within and across participants (**Fig. S4**).

Neurons in supramarginal gyrus (SMG) in two of the three participants (S2, S3) showed robust encoding of grasp type during motor planning (mean latency: 425 ms, accuracy: 60%) and imagined execution (mean latency: 511 ms, accuracy: 67%) (**Fig. 2C,D; see Methods**), indicating sustained representation of motor engagement, independent of object presence or object type (**Fig. 3J,K**). Participant S4’s SMG implant showed little neural representation of grasp or object properties; notably, the implant location was also anatomically distinct from S2 and S3 (**Fig 1A**). Positioned far more medially, on the lateral bank of the intra-parietal sulcus, our data suggests implantation occurred in a different sub-region of SMG (see Discussion).

Recordings from the medial gyrus of the anterior intraparietal cortex (AIP) also demonstrated encoding of grasp type during motor planning (mean latency: 541 ms, accuracy: 45%) and imagined execution (mean latency: 589 ms, accuracy: 58%) (**Fig. 2C,D**). During motor planning, AIP encoded grasp type (**Fig. 2D**) and object shape (**Fig. 2G**). During imagined execution, AIP encoded only grasp type (**Fig. 2D**).

Populations of neurons in the motor cortex (MC) robustly represented grasp type throughout both the planning (mean latency: 383 ms, accuracy: 69%) and imagined execution (mean latency: 211 ms, accuracy: 85%) phases of the task (**Fig. 2A,B,C**). Across both participants (S3, S4), decoded grasp type was significantly higher than any other individual area (**Fig. 2B**), even during the planning phase. While robust grasp representation during motor planning was unexpected, our group has selectively observed planning activity in MC, only when the region is not actively engaged in a motor task^29^. While our data suggests concurrent planning activity across MC and PPC, motor cortex leads the onset of neural activity during imagined execution (**Fig. 2C**).

Across participants, somatosensory cortical neurons represented grasp type during imagined execution (**Fig. 2D**), but only neurons within the receptive field of the hand (outside accuracy (S2): 26% vs. inside accuracy (averaged S3 and S4): 64%; chance: 25%). Participant S2’s sensory arrays (S1) are in the receptive field of the arm^30^ (outside the receptive field of the hand) and thus showed little representation of grasp type (**Fig. 2A,B**: S2). The time course of grasp representation in S1 roughly mirrors AIP (**Fig. 2A,B**: S3, S4), suggesting efference copies (or projections) from posterior parietal and/or precentral gyrus may be driving this activity when implants are within the task-relevant receptive field^31^.

### Cognitive processes influence unit selectivity in each region

Single-unit tuning to grasp occurred in both Planning and Action phases in all CGN regions (**Fig. 3-5C,D,H**). Tuned units could be selective (tuned to only one grasp type) or mixed-selective (tuned to two or more grasp types). A predominant pattern emerged in all CGN regions where during motor planning, the proportion of selective to mixed-selective units was roughly equivalent or favored selectivity (**Fig. 2E**). However, during imagined execution, there was a higher proportion of mixed-selective units. This finding was largely consistent across participants and cue contexts (**Fig. S5**). This pattern may reflect that during motor planning, receiving visual feedback of the exact grasp type the participant intends to perform could lead to activating a more selective population; whereas without any feedback, the populations are more broadly activated and thus mixed-selective. SMG was an outlier in that the population was roughly equivalent or favored selectivity during both phases.

### SMG demonstrates partial mixed selectivity at the population level

Supramarginal gyrus encodes grasp type, object size, and object shape from the single-unit to the population level, suggestive of partial mixed selectivity encoding within the neural network.

Populations of neurons in SMG significantly encoded grasp throughout motor planning and imagined execution (**Fig. 3A**; Wandelt et al. (2022)^20^). Complete information about grasp type appeared early during planning (peak latency: 681 ms, accuracy: 67%) and imagined execution (peak latency: 851 ms, accuracy: 60%), reaching peak decoding accuracy before any other region (**Fig. S6**). However, information about grasp type extended beyond population-level activity.

Encoding of grasp was evident at the single-unit level, with many single-units responding selectively and uniformly to grasp types during both motor planning and imagined execution (**Fig. 3B,C**).

Although grasp types were highly separable in SMG neural populations, distinct representations were found between planning and imagined execution. Utilizing a cross-phase decoding analysis (**see Methods**), we compared representations using a grasp linear-decoder trained on one phase (i.e. Planning) but tested on another (i.e. Action). Grasp encoding *completely generalized* from planning to imagined execution, with no significant difference between decoders trained during either Planning or Action and tested on Action (**Fig. 3I**). However, *incomplete generalization* was observed from imagined execution to planning (decoders trained on Action and tested on Planning were significantly lower than those trained on Action). While many tuned units are shared between phases (S3: 53%, S2: 30%), the largest proportion of units tuned only to grasp, occur during imagined execution (S3: 34%, S2: 23%) (**Fig. 3H**). This data suggests additional neural recruitment or distinct neural processes occurring solely during imagined execution (**Fig. 3I**), largely consistent across contexts and participants (**Fig. S8, S9**).

### Grasp encoding is context-dependent in SMG

In our previous work, grasp encoding in SMG was demonstrated using images depicting a hand interacting with an object (Wandelt et al., 2022). We hypothesized by separating the image into its individual components might result in contextually specific encodings.

We found grasp encoding was modulated by the context in which a grasp was presented, while still retaining shared grasp-related structure across contexts. Cross-context decoding (**see Methods**) showed that grasp representation was partially context-specific; while cross-decoding accuracies generally remained above chance, decoding accuracies significantly varied between contexts, demonstrating incomplete generalization among contexts (**Fig. 3E**). Context-specificity was particularly evident during planning but remained present in the Hand and Hand+Object contexts during imagined execution as well. Further, contexts involving a hand generally produced high classification accuracies during motor planning and imagined execution (**Fig. S10**). A higher proportion of units were tuned to grasp in SMG during planning when both the hand and object were presented compared to the hand or object presented alone (**Fig. 3D**). From these tuned units, distinct populations were recruited within each context particularly during planning (Hand+Object: 28-30%, Hand only: 13-24%, Object only: 6-18%) (**Fig. 3F,G**). We also observed overlap among intuitive contexts (7-22% Hand+Object and Hand only, compared to 2- 3% Hand only and Object only). These context-specific units may be driving the increase in classification accuracy for contexts involving the hand and the incomplete generalization among contexts.

### SMG encodes object properties

Due to SMG’s role in tool use and grasping, we hypothesized the object properties themselves would also be encoded. Across all grasp types, single units tuned to size of the object (Small vs. Large) (**Fig. 3L**) and object shape (**Fig. 3N**) were identified in SMG. Distinct subpopulations of size- (**Fig. 3M**) and shape-specific units (**Fig. 3O**) arose, suggesting the encoding of object properties at the single-unit level. At the population level, both size (**Fig. 3P**) and shape (**Fig. 3Q**) were significantly decoded during planning and persist into imagined execution.

### Grasp encoding in SMG remains stable when manipulating object properties

When manipulating the size and the shape of the object being grasped, grasp decoding remained stable (**Fig. 3J,K**), suggestive of unique neural sub-populations, each encoding grasp and object independently (**Fig. 3M,O**).

### AIP encodes selective object properties

AIP demonstrates selective encoding, such that grasp and some object properties are encoded. Consistent with findings from NHP AIP, object shape was well represented in human AIP; however, object size was not (**Fig. 4P**).

Throughout motor planning and imagined execution, populations of neurons in AIP significantly encoded grasp type (**Fig. 4A**). Even though this area is predominantly thought of as a planning region, for one participant (S4), grasp encoding was much stronger during imagined execution and was markedly similar to the primary sensorimotor areas (**Fig. S11**, see Discussion). One possible explanation for this discrepancy is strongly maintained connections within the CGN due to increased intact sensation relative to our other participant, and therefore a stronger similarity in AIP performance to MC and S1. Encoding of grasp was also seen at the single-unit level, with units selectively and uniformly tuned to grasp during motor planning and imagined execution (**Fig. 4B,C**).

AIP exhibits strong phase-dependent grasp encoding during motor planning versus imagined execution, with incomplete generalization between the phases (**Fig. 4I**), consistent across contexts and participants (**Fig. S9**). This may represent a more active integration process in AIP during planning before receiving feedback/input from the sensorimotor loop during imagined execution. Across participants, 33-41% of units were shared between both phases and 33-55% were uniquely tuned during Action, further suggesting unique processes during motor planning and imagined execution (**Fig. 4H**).

### Grasp encoding is context-dependent in AIP

Grasp encoding was also modulated by context. During both planning and imagined execution, decoding predominantly either incompletely generalized or did not generalize at all among contexts (**Fig. 4E**). This finding may be partially explained by a different proportion of units recruited during each context, with Object only recruiting fewer units during planning (**Fig. 4D**). However, although a similar proportion of units were tuned to grasp across the three contexts during imagined execution (Fig. 4D) and we observed a large amount of overlap among all contexts during imagined execution (19-51%) (**Fig. 4F,G**), the cross-context decoding suggests that these units did not encode grasps in a fully context-invariant manner (**Fig. 4E**). Additionally, we observed a large, unique population of grasp-tuned units when the hand and object were presented together during planning (17-49%), further supporting that context influences grasp encoding in AIP.

### Grasp encoding in AIP remains stable during planning when manipulating object properties

When altering object size and shape, grasp type could still be significantly decoded during motor planning (**Fig. 4J,K**). Grasp type decoding remained stable throughout the trial when altering size. Due to natural degradation of the micro-electrode arrays^32^, S4’s AIP data was excluded from analyses related to size and shape (see Discussion; **Fig. S12**).

### AIP selectively encodes object properties

Object shape was encoded in this region of human AIP, but not object size. Many neurons in AIP demonstrate tuning to object shape predominantly during motor planning, with proportions that rival grasp-tuned units (**Fig. 4N,O**). Populations of neurons significantly decoded object shape during planning, at accuracies comparable to grasp (**Fig. 4Q**). In contrast, while there were units tuned to object size (**Fig. 4L,M**), there was no significant decoding of size throughout the trial (**Fig. 4P**).

### MC encodes motor-related variables

Motor cortex (MC) neural populations encoded grasp type and object size, but not object shape. Prior literature supports MC’s role in encoding motor variables^5^, but MC’s role in encoding object properties like shape in NHPs^5,33^ remains unanswered in humans. Our results suggest object “size” was well represented (**Fig. 5P**), but not shape (**Fig. 5Q**). In our task, object size could be correlated with hand aperture, and given MC’s prior known role in representing motor variables, we will refer to object size as hand aperture throughout this section (see Discussion).

Populations of neurons in MC significantly encoded grasp type throughout the trial, with robust classification and tuning during planning and imagined execution (**Fig. 5A,D**). MC had the highest grasp classification of all CGN regions both at the timebin-level and when averaged across entire phases (**Fig. 2A,B**, green line). Many single units in MC selectively encoded grasp type and these units were uniformly distributed across grasp types (**Fig. 5B,C**).

Even in MC, motor planning and imagined execution largely demonstrated incomplete generalization during cross-phase decoding, as evidenced by significantly higher grasp classification of the training phase than the testing phase during planning (S3) and imagined execution (S3, S4) (**Fig. 5I**). This finding was largely consistent across contexts and participants (**Fig. S9**) and suggests partly distinct neural processes during both phases. This result was consistent with the composition of grasp-tuned units during Planning and Action phases. Most units were either tuned during both phases or tuned only during imagined execution, with only 3- 4% uniquely tuned during motor planning (**Fig. 5H**).

### Strongest grasp encoding in MC results when contexts involve a hand

When we examine the subpopulations of neurons tuned to grasp in each context, the story becomes more complex. Despite stable single neuron representation across planning and imagined execution (**Fig. 5F,G**), the population decoding was dominated by hand representation (93-94% Hand peak accuracy vs. 73-82% Object peak accuracy, **Fig. S10**). Hand and Hand+Object contexts demonstrated complete generalization, suggesting a shared grasp representation that incompletely generalizes to the Object only context (**Fig. 5E**). While the proportion of neurons encoding grasp during the Object context were similar among contexts, a reduced total number of neurons may partially explain this result (**Fig. 5D**).

### MC encodes hand aperture

MC encoded grasp and grasp-relevant information (hand aperture), but not the object property of shape, in contrast to recent work in NHPs^33^ (see Discussion). As expected, individual neurons responded selectively to hand aperture predominantly during imagined execution (**Fig. 5L,M**).

At the population level, hand aperture was significantly decoded as early as the planning period, but most strongly and sustained during imagined execution (**Fig. 5P**). As in SMG and AIP, grasp decoding in MC remained stable when manipulating object properties (**Fig. 5J,K**).

Interestingly, while we observed the occasional single neuron modulated by object shape, these units were rare and predominantly also tuned to grasp (**Fig. 5N,O**). In conjunction with no significant decoding of object shape, our data suggests MC did not encode object properties (**Fig. 5Q**).

## DISCUSSION

Throughout the human cortical grasping network (CGN), grasp and object properties were robustly represented both at the single neuron and population level, during motor planning and imagined execution. Across three (*right-handed*) human participants, chronically implanted in the contra-lateral hemisphere (**Fig. 1A**), grasp was significantly represented in each region of the cortical grasping network (**Fig. 2A**), including supramarginal gyrus (SMG), anterior intraparietal cortex (AIP), and the hand-knob region of the motor cortex (MC). Somatosensory cortex (S1) also encoded grasp during imagined execution, within the receptive field of the digits (**Fig. 2D**).

Our results suggest different processes occur during motor planning and imagined grasping. While grasp representation increased during imagined execution for most regions (**Fig. 2A**), object properties were often more strongly encoded during planning (**Fig. 2G**). Activity related to planning activates the entire network simultaneously (ns, p = 0.5 (S3), p = 0.625 (S4))^34^.

During imagined execution, MC appeared to activate before all other regions, though there were no significant differences among regions (**Fig. 2C**). Additionally, even pre-frontal cortex (dlPFC) responded to motor engagement, suggesting full activation of the fronto-parietal network (**Fig. 2C**). Surprisingly, the hand knob region of MC strongly encoded planning information, adding further evidence that the crown of the pre-central gyrus may be included in the hypothesized gradient from primary motor to pre-motor regions (Fig. 4F of Ariani et al., 2025)^35^. Grasp was encoded throughout the CGN and PPC regions strongly encoded object properties during both planning and imagined execution (**Fig. 2F,G**). Motor cortex also strongly encoded correlates of object size (likely hand aperture) during imagined execution, but not object shape (**Fig 2F**).

### Context matters

Developing a robust neuroprostethic assistive device for performing dexterous manipulations of objects and activities of daily living (ADLs) will require demonstrations of flexible, context-independent control in real-world naturalistic environments. An important component in achieving these goals is the development of accurate BCI algorithms, which quickly translate recorded cortical signals of the participant’s motor intention into action^36^. Previous work has shown many context-dependent effects on control signals from MC, including object interaction^27,37,38^, immersive environment^39^, cognitive load^40^, arm posture^41^, task goals^42^, reward^24^, concurrent speech^43^, and sensory feedback^44,45^. To achieve stable performance, either all these effects must be accounted for or cortical signals independent of context must be identified.

In our work, we observed regions during a variety of contexts (planning vs. execution) and object interaction (a hand and object together vs. its individual components), and found both affect the neural representation of grasp. We hypothesized that different contexts could result in distinct neural control strategies. Visual contexts with an explicit hand position (Hand only) might engage a position-based encoding of individual digit endpoint placement, whereas object interaction contexts (Hand+Object) might activate mechanisms associated with grasp stabilization or object squeezing (isometric force production). Additionally, viewing the object by itself (Object only) is typically how humans encounter items in their daily lives.

Understanding how context influences our ability to classify grasp can help us determine how best to train and utilize decoders for a grasp BMI.

Our results demonstrate the neural processes occurring during motor planning and imagined execution are partly distinct in each brain region (**Fig. 3I,4I,5I**). Specifically, these results suggest it may be possible to design BMI decoders which selectively engage during imagined execution, thereby reducing the likelihood of premature grasp execution. This is important in BMIs because it affords the BMI user agency over when to perform the movement. Additionally, as demonstrated with other motor modalities (i.e., reaching)^46^, our participants can perform highly accurate grasp with purely imagined activity. As shown with motor control of speech, imagined actions may be even easier to perform and produce less fatigue for the user than attempted actions^47^.

Across all regions and participants, when the object was presented alone (O only), we observed lower grasp classification accuracies compared to contexts where the hand was presented (H, or H+O) (**Fig. S10**, yellow bar). Despite the same verbal task instructions, different visual cues shown during the planning phase (hand alone, hand+object, and object alone) were significant enough to impact representation during imagined motor execution (**Fig. 3E,4E,5E**). There are a few possible explanations. Firstly, it is possible different networks may be recruited when a hand is explicitly shown^48,49^. In our data, we observed different contexts recruiting different subpopulations in each region, though many units were tuned to grasp in all three contexts (**Fig. 3-5F,G**). Secondly, while participants were instructed to perform the same action, it is possible that each participant may have a unique strategy, in addition to potential differing motor imagery strategies across contexts. Strategy variance between contexts could affect decoder performance^50^. Thirdly, lack of an explicit hand position (Object only) during planning could also lead to increased variance of participant’s imagined grasp. Fourthly, neurons encoding the relative position between the hand and the object, the reach vector, which has been shown in PPC and premotor cortex of NHPs^51,52^, could be absent when the object is presented alone and decrease decoder performance.

The ability to decode object information could be incredibly useful, as evidenced by the presence of objects severely affecting the ability to operate grasp BMIs^27^. The fact that we could decode all three variables in PPC (grasp, size, and shape), paired with fast latencies and high classification accuracies, make PPC a desirable implant location for grasp BMIs.

### Grasp type can influence object property decoding in SMG

Our data exhibited partial mixed selectivity within SMG at the population level, encoding grasp type, object size, and object shape, suggesting a crucial role in processing motor plans and object characteristics. SMG demonstrated a “preference” for specific grasp types over others when classifying object size. For MediumWrap and Sphere3Finger grasp types^53^, we could decode size with higher accuracy during both motor planning and imagined execution compared to Lateral and PalmarPinch grasp types (**Fig. S13A**). If SMG encodes positional information of the intended end effectors, the required increased dimensionality of the control signal with increased number of effectors (i.e. the virtual finger hypothesis^54^), may increase the separability of the population activity. This preference was not observed across other areas (**Fig. S13B**); thus, this finding is unlikely due to increased task demands or complexity of the particular grasp conditions.

### Human AIP may be distinct from NHP AIP

Our results demonstrate that neurons in the medial gyrus of the anterior intraparietal cortex (human AIP) selectively encoded object characteristics (i.e. shape) (**Fig. 4N,O,Q**), compared to the oft-referenced NHP AIP homologue, shown to predominantly encode object properties (i.e. size and shape^5^).

Size was not significantly decodable during motor planning in AIP (**Fig. 4P**). This was surprising, given historical evidence for involvement in planning or higher-order processing^8,13,16^ and NHP studies have demonstrated homologue AIP plays a strong role in object processing^5,7,55^. It is possible that the human cortex has multiple cortical regions dedicated to object representation and manipulation, rather than centralized in AIP. SMG is a likely candidate given our data demonstrated representation of object properties in SMG, imaging data has suggested SMG’s role in tool use^18,56^, and there is no known homologous SMG in NHPs. Support for this hypothesis includes enhanced human hand dexterity driven by complex object/tool use^57,58^, structural reorganization of association cortices throughout evolution^59^, human-specific multi-region lateralized networks that allow fronto-parietal connectivity^60^, connectivity driving neural dynamics^61^, and the postulated connectivity between AIP and SMG^18^.

### AIP populations are selectively activated by context

Our data suggests variable and context-dependent encoding of grasp in AIP across two participants. In one participant, we observed highly selective tuning for object and hand interaction (**Fig 4D**, S3). During motor planning, we observed the highest percentage of grasp- tuned units when the hand and the object were interacting, the lowest percentage when the object was presented alone, and the hand presented alone falling between them. While this result is unlikely to be explained solely due to increased complexity of visual stimuli (**Fig. S14B**), our data may be further evidence of compositional encoding within AIP^62^. With approximately 20% of the neurons in both phases shared among the contexts (**Fig. 4F,G**), each participant had significant populations of uniquely tuned neurons to each context. These three contexts may also selectively activate known sub-populations of AIP neurons, preferential to encoding visual information (corresponding to the object alone), motor information (hand alone), or a combination known as visual-motor (hand and object together)^55^.

### Planning activity present in MC

While NHP literature is mixed on MC’s role in object processing^5,33^, our results demonstrated human hand knob MC encoded motor-related variables, such as grasp type (**Fig. 5A,B,C**) and hand aperture (**Fig. 5L,M,P**) during motor planning and imagined execution. When MC is not actively engaged in a motor task, it is possible that it may encode information from other connected regions (including PPC). This has been previously demonstrated in our lab, with motor cortex encoding the next movement plan unless it is actively engaged in a motor task, whereby the plan encoding disappears^29^. This has also been previously observed in primary motor cortex during bimanual movement^63^. When only the ipsilateral hand is engaged, the contralateral motor cortex encodes the movement. When both hands are engaged, the contralateral cortex suppresses activity from the ipsilateral side. Additionally, complex movements such as grasp demonstrate correlated contralateral/ipsilateral representations compared to more independent representations for reach^64^.

### MC encodes hand aperture rather than object size

MC significantly encoded hand aperture during motor imagery. While it is possible that MC encoded an object property such as size, we believe hand aperture was encoded rather than object size. First, we could predominantly decode this variable during imagined execution rather than motor planning (**Fig. 5P**), with the significant size decoding during motor planning occurring when the hand alone was shown. This suggests involvement in motor commands rather than object processing, which we would expect to see during motor planning, as we did with SMG and AIP. Second, the “preferred” grasp types with the highest size classification accuracies are the ones with the largest delta for hand aperture (**Fig. S13B**). Anecdotally, participant S4 reported “feeling the stretch” in their hand when imagining performing these grasps, and size or hand aperture was decodable from S1 during movement (**Fig. 2F**). Third, the literature overwhelmingly supports MC’s involvement in processing motor variables^5,7,9,10^ like hand aperture, rather than object properties, like object size. Further, Schaffelhoefer & Scherberger (2016) found that NHP MC (M1) only encodes motor information (not object size or shape) and that this only occurs during imagined execution^5^. One recent study in NHPs did find object property encoding in MC^33^. Our results may differ from this finding as their study was done in fully intact non-human primates, whereas our participants are humans with tetraplegia due to spinal cord injury. Additionally, their study utilized dynamic grasping of objects, whereas our participants used motor imagery cued by static images.

### MC is modulated by object properties

While we see MC modulated by object properties during both motor planning and imagined execution, these properties are not modulating MC in such a way that they are encoded in this region. Rather, it seems these properties affect the firing rates resulting in tuning (**Fig. 5O**), but not in a way that is unique, and therefore classifiable, for each object shape (**Fig. 5Q**). This is similar to the finding in Downey et al. (2017) which showed the presence of an object alters MC (M1) firing rates^27^.

### Grasp is encoded in S1 during imagined grasping

We observed significant classification of grasp type from S1 during imagined execution in two out of three participants (**Fig. 2A,B**). We believe this discrepancy among participants is due to the location of their S1 implants. Participant S2’s S1 implants were located in the receptive field of the arm, while participants S3’s and S4’s S1 implants were in the receptive field of the hand. In a previous study from our lab, S1 encoded reach direction during an imagined reaching task with participant S2^31^. As this task engaged imagined movement of the arm, it follows that S2’s S1 arrays demonstrated encoding of reach direction. However, because our tasks relied on imagined hand movements, we did not observe significant classification related to grasp in participant S2, but we did in participants S3 and S4. This suggests that imagined movements can result in movement encoding in the primary somatosensory cortex if recording from within the receptive field of the movement (**Fig. S7D, S8D, S9D**).

### Results are due to grasp and object encoding, not visual stimuli

One may reasonably consider that the neural tuning differences we see among the three contexts (Hand only, Object only, Hand+Object) are the result of differing visual statistics, with the object containing the least amount of visual information and the object presented with the hand containing the most. To test this hypothesis, we performed the Pixelated Task by interspersing trials where participants are cued normally (“Regular”) or cued with pixelated images of each grasp/object (“Pixelated”) (**Fig. 1E**; see Methods for details). A very small number of units were tuned to pure visual stimuli during the motor planning, but this amount was not significantly different among contexts and did not significantly contribute to the total number of tuned units that were recruited by Regular images (**Fig. S14A**). This suggests any differences among contexts were a result of differences in how the hand, object, and their interaction affected motor planning rather than differences in visual statistics presented.

### Task relevant participant performance

Participants correctly performed the imagined execution during the Action phase of the task. Results from No-Go trials demonstrate equivalent percentage of tuned units throughout the trial until the Action phase, where there was a sharp drop-off of tuned units to baseline (ITI) levels quickly after the presentation of the No-Go cue (**Fig. S14B**). This indicates the participants were imagining executing the grasp only during Go trials, and that the tuned units we observed during the Go trials were directly related to performing the imagined execution.

### Subject Differences

Across the three SMG implants, participant S4’s did not significantly encode grasp, size, or shape, in sharp contrast to our other two participants (**Fig. 2A**, S4), despite maintaining similar numbers of total recorded neurons (**Fig. S4**). SMG is hypothesized to contain different specialized sub-divisions related to language, phonetic processing, tool use, visual and sensory integration among others^65,66^. We did find motor task modulating neurons, but little unique selectivity in individual grasp types (**Fig. S12A**), suggesting this implant was in a different sub- region of SMG than the other two participants. Additionally, the neural population from S4’s SMG implant could not significantly decode object size or object shape (**Fig. S12B**). A difference in anatomical location of S4’s SMG implant relative to S3’s and S2’s, may also support our hypothesis of a different SMG sub-region (**Fig. 1A**).

With S4’s AIP array, we obtained much higher grasp classification accuracies during imagined execution compared to motor planning (**Fig. 4A**, dotted line). This was surprising, as we consider AIP to be a higher order region involved in movement planning. Interestingly, between AIP and S1, when controlling for equal neural population size, decoding performance was identical (**Fig. 2B**), suggesting this subregion of PPC may be receiving efferent copies from MC, like S1, resulting in similar results among the three regions. Additionally, this region demonstrated the same results as primary sensorimotor regions during cross-phase and cross-context decoding (**Fig. S11**). Due to typical degradation of S4’s AIP array, fewer neurons were recorded between the Go/No-Go Task (grasp) and the Varied Sizes Task (size) and subsequently the Disassociation Task (shape). This is apparent in loss of ability to classify grasp and chance level classification of the object properties (**Fig. S12C**).

## CONCLUSIONS

In this study, we elucidated the roles of key regions within the human cortical grasping network using single-unit firing rates from populations in SMG, AIP, and MC during motor imagery. AIP plays a role in grasp and object shape processing during motor planning, SMG works to encode grasp and object properties, and MC functions to both plan and execute imagined motor actions. Mixed-selectivity encoding in SMG could make it an optimal target for grasp BMIs, by decoding grasp while also incorporating information about the object.

## RESOURCE AVAILABILITY

### Lead contact

Requests for further information and resources should be directed to and will be fulfilled by the lead contact, Mackenzie Thurston.

### Materials availability

• This study did not generate new unique reagents.

### Data and code availability

• All data produced in the present study will be publicly available as of the date of publication.

• All original code produced in the present study will be publicly available as of the date of publication.

• Any additional information required to reanalyze the data reported in the paper is available from the lead contact upon request.

## ACKNOWLEDGMENTS

We would like to thank our participants, FG, AN, and GB, for their hard work and engagement in this project; without you, this research would not be possible. This research was supported by NIH NINDS U01NS123127, the James G. Boswell Foundation, Craig H. Neilsen Foundation, and the T&C Chen Brain-Machine Interface Center. A special thank you goes to B. Liu, B. Ruzsala, S. Darcy, N. Mynhier, L. Lin, and S. Lee for their insightful discussions.

## AUTHOR CONTRIBUTIONS

Conceptualization, M.J.T., R.A.A., D.B., and S.K.W.; methodology, M.J.T., D.B.; Investigation, M.J.T.; writing—original draft, M.J.T.; writing—review & editing, R.A.A., D.B., S.K.W.; funding acquisition, R.A.A., K.P., and D.B.; resources, R.A.A. and K.P.; Clinical Trial Management, K.P., R.A.A., D.B.; supervision, R.A.A. and D.B.; Surgical and Medical Intervention, B.L. and C.L.

## DECLARATION OF GENERATIVE AI AND AI-ASSISTED TECHNOLOGIES

During the preparation of this work, the author(s) used ChatGPT to verify and condense code used for data analysis. After using this tool or service, the author(s) edited the code as needed and take(s) full responsibility for the content of the publication.

## ADDITIONAL RESOURCES

This research is part of a human brain-machine interface clinical trial (NCT01964261): https://clinicaltrials.gov/study/NCT01964261?id=NCT01964261&rank=1

## SUPPLEMENTAL INFORMATION

Supplementary Figures S1–S14

## METHODS

### Experimental Model and Subject Details

This project was completed with three tetraplegic participants that were recruited for an IRB- and FDA-approved clinical trials of a brain-machine interface. All participants gave informed consent to participate. Participant S2 has a spinal cord injury (SCI) at cervical level C5 and was injured 2 years prior to enrollment. Participant S3 has a SCI at cervical level C5-6 and was injured 3 years prior to enrollment. Participant S4 has a SCI at cervical level C5 and was injured 51 years prior to enrollment.

### Implants

The targeted areas for implantation across participants were the left supramarginal gyrus (SMG), anterior intraparietal cortex (AIP), motor cortex (MC), primary somatosensory cortex (S1), ventral premotor cortex (PMv), and dorsolateral prefrontal cortex (dlPFC). S2 has four arrays, one 96- channel multi-electrode array (Neuroport Array, Blackrock Microsystems, Salt Lake City, UT) in SMG and PMv and two 7 x 7 sputtered iridium oxide film (SIROF)-tipped microelectrode arrays with 48 connected channels each in S1. S3 has 6 multi-electrode array implants (81 electrode SIROF array, with 64 connected, Blackrock Microsystems, Salt Lake City, UT) in SMG, AIP, MC, PMv, and 2 arrays in S1. S4 has 6 multi-electrode array implants (81 electrode SIROF array, with 64 connected, Blackrock Microsystems, Salt Lake City, UT), two in S1, one in SMG, AIP, MC, and dlPFC. Exact implant sites for each participant were determined using fMRI as described in Aflalo et al. (2015)^51^. This work focused on the SMG, AIP, MC, and S1 implants.

### Data Collection

Single-unit neural data for this work was collected between 2023 and 2025. We recorded broadband electrical activity from the arrays using Neural Signal Processors developed by Blackrock Microsystems and Ripple Neuromed interchangeably from August 2023 to May 2025. Analog signals were amplified, bandpass filtered (0.3-7500 Hz), and digitized at 30,000 samples/sec. Data were bandpass filtered again (250-5000 Hz) with a threshold at -4.5 the estimated root-mean-square voltage of the noise to identify action potentials. Channels were included from a session in post-processing by achieving a minimum firing rate of 2 Hz. Channels that do not meet this threshold are excluded from analysis. Multi-unit activity at each channel, in spikes/sec (Hz), are the data underlying this work.

#### Breakdown of number of recorded units per session

*Breakdown of number of recorded units per session:*
| Participant | Altered Context Task | Varied Sizes Task | Disassociation Task | Pixelated Task |
| --- | --- | --- | --- | --- |
| **S2** | SMG: 30 ± 3<br>PMv: 10 ± 3<br>S1: 52 ± 2 | SMG: 48 ± 7<br>PMv: 23 ± 6<br>S1: 59 ± 6 | SMG: 54 ± 11<br>PMv: 41 ± 11<br>S1: 65 ± 9 | SMG: 27 ± 3<br>PMv: 23 ± 11<br>S1: 57 ± 8 |
| **S3** | SMG: 62 ± 0<br>PMv: 36 ± 2 | SMG: 62 ± 1<br>PMv: 38 ± 1 | SMG: 61 ± 1<br>PMv: 35 ± 2 | SMG: 62 ± 1<br>PMv: 43 ± 5 |
|  | S1: 50 ± 11<br>AIP: 44 ± 2<br>MC: 63 ± 0 | S1: 90 ± 6<br>AIP: 46 ± 1<br>MC: 61 ± 1 | S1: 98 ± 6<br>AIP: 48 ± 1<br>MC: 61 ± 1 | S1: 70 ± 6<br>AIP: 49 ± 2<br>MC: 62 ± 1 |
| **S4** | SMG: 61 ± 1<br>S1: 108 ± 6<br>AIP: 60 ± 1<br>MC: 61 ± 1<br>dlPFC: 25 ± 1 | SMG: 42 ± 3<br>S1: 70 ± 2<br>AIP: 34 ± 2<br>MC: 52 ± 1<br>dlPFC: 18 ± 1 | SMG: 44 ± 3<br>S1: 64 ± 6<br>AIP: 25 ± 6<br>MC: 51 ± 1<br>dlPFC: 18 ± 1 | N/A |

### Experimental Tasks

We implemented a motor imagery task that cued four different grasp types with static visual images originally selected from the “Human Grasping Database” (Feix et al., 2016)^28^ and modified using Unity. The grasps were selected because they represent some of the most common grasps used during activities of daily living (ADLs) and cover the range of grasp taxonomies. They are labeled “Lateral,” “MediumWrap,” “PalmarPinch,” and “Sphere3Finger” (**Fig. 1C**).

#### Go/No-Go Task

Each trial consisted of four phases: ITI, Planning, Delay, and Action (**Fig. 1B**). Trials were performed in blocks of three contexts: a hand alone (H), a hand grasping an object (HO), or an object alone (O). Each trial began with a 2-second intertrial interval (ITI), in which the screen was blank, and the participant was asked to clear their mind. Then participants were visually cued with a static image of one of four grasp types (**Fig. S1A** for all variations). There was a brief 0.5 second delay before the Action phase, where the participant saw either a green or red dot for 4 seconds. The green dot cued the participant to either imagine performing the grasp that was shown (H) or imagine performing the grasp and interacting with that object (HO and O). The red dot cued the participant to abort the motor plan and perform no imagined action.

#### Varied Sizes Task

A variation of the task of the <u>Go/No-Go Task</u> was designed, and the overall trial structure was maintained. We altered each hand to create the smallest and largest feasible grasps for each grasp type (**Fig. 1D; Fig. S1B** for all variations). During the Action phase, participants were given the same instructions to perform imagined motor imagery, paying particular attention to adjusting the size of the grasp accordingly. During blocks where only the object was presented (Object only), small blue dashes were added to indicate which fingers were to be used and where they were to be placed.

#### Disassociation Task

Another variation of the task was created where we maintained the same trial structure as the <u>Go/No-Go Task</u>, but during Planning, a static image of one grasp type interacting with one object was the stimulus every time. This stimulus varied, as we paired each grasp type with each object shape, such that each object was grasped by all 4 grasp types (**Fig. 1E; Fig. S1C** for all combinations). Dimensions of each shape were altered slightly to ensure feasible grasping by each grasp type. This task variation had no hand only or object only contexts, as the grasp and object pairs are now decoupled and thus a hand and object must always be used to cue which grasp/object combination to perform. The hand configuration was held constant for pairing with each object (i.e., the Lateral grasp appeared the same no matter what object it was going to grasp). Similarly, the object shape was held constant across pairings with each grasp type. As some grasps would have obscured the object and introduced a visual confound, each object was depicted as slightly in front of the hand (prior to interaction).

#### Pixelated Task

A visual control task was created where we pixelated the images previously presented in the <u>Go/No-Go Task</u> (**Fig. 1F; Fig. S1D** for all variations), such that the grasps and objects were unrecognizable to the participants. Half of the trials followed the exact trial structure outlined above, and half of the trials consisted of pixelated versions of those same images. Images were presented in random order. Participants received instructions to visually take in the pixelated image during the Planning phase and perform no imagined motor action during the Action phase. Pixelated images share the same visual statistics as Regular images without being recognizable as a specific grasp type or object shape.

#### 50:50 Task

The same task as the original Go/No-Go Task was performed, but the Go and No-Go conditions were performed at equal rates to ensure adequate comparison.

#### Breakdown of number of sessions

***Breakdown of number of sessions:***
| Subject | Go/No-Go Task | Varied Sizes Task | Disassociation Task | Pixelated Control Task | 50:50 Control Task |
| --- | --- | --- | --- | --- | --- |
| S2 | 7 | 8 | 8 | 9 | 0 |
| S3 | 6 | 8 | 6 | 6 | 5 |
| S4 | 7 | 7 | 7 | 0 | 0 |

#### Neural firing rates

Units that crossed the threshold (ie., spiked) during any trial were included in analysis. Instantaneous firing rates (spikes/sec) of these units were computed as the number of spikes that occurred in 50ms bins, divided by 50ms. This value was then smoothed using a Gaussian filter with a kernel width of 50ms. For the Go/No-Go Task, 40 trials were recorded per block (8 repetitions of 4 grasps = Go trials, 2 repetitions of 4 grasps = No-Go trials), two blocks per context (HO, H, and O) every session. For the Varied Sizes Task, 28 trials were recorded per block (2 repetitions of 4 grasps x 2 sizes = Go trials, 1 repetition of 4 grasps x 2 sizes = No-Go trials), two blocks per context every session. For the Disassociation task, 40 trials were recorded per block (2 repetitions of 4 grasps x 4 objects = Go trials, 2 repetitions x 4 grasps = No-Go trials), with three blocks recorded every session, along with one block of each context (HO, H, and O) for direct comparison. For the Pixelated Task, 24 trials were recorded per block (3 repetitions x 4 grasps = Regular images, 3 repetitions x 4 pixelated grasps = Pixelated images), two blocks per context every session. For the 50:50 Go:No-Go Task, 32 trials were recorded per block (4 repetitions of 4 grasps for Go and No-Go trials), two blocks per context every session. Depending on the task, the number of trials were adjusted for the comfort of the participant to 20-28 trials per block to accommodate Participant S2 with shorter tasks.

For each Task, trials where the participant physically moved, experienced a disturbance (e.g., received a phone call or was clearly not actively engaged in the trial) were removed from data analysis.

## QUANTIFICATION AND STATISTICAL ANALYSIS

All analyses were performed using MATLAB R2023b.

### Single-unit identification

In this work, unit is used to refer to the activity recorded from a single microelectrode. Threshold crossings were binned in 50 ms timebins and converted into instantaneous firing rates, as described above. To identify single-unit responses, the average firing rate (Hz) per timebin of each channel was calculated as a function of each feature (grasp type, object shape, size) within sessions. 95% confidence intervals were calculated by bootstrapping 1000 times and calculating the standard error of the mean (SEM).

### Linear regression analysis

Units that selectively fire (tuned) for the different variables were identified by linear regression in two ways: 1) step by step in 50 ms timebins to assess for tuning across the entire trial; 2) average firing rates across the Cue (2s) and Action (4s) phases to compare between both phases. The model returns a fit that estimates the firing rate of a unit based on the following variables:

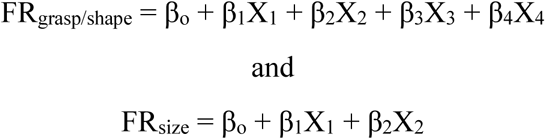

where FR represents the firing rate of that unit and β corresponds to the estimated regression coefficients. X is a one-hot encoding indicator variable that contains which data correspond to which feature (grasp type, object shape, size). The first 8 rows of X correspond to the ITI phase and thus only contain zeros. The baseline variable, βo, is the average firing rate during the ITI phase. The other rows of the indicator variable indicate the trial data, such that each row contained one numerical value (1) that corresponds to a feature (grasp types 1-4, object shapes 1- 4, sizes 1-2). For example, if the first trial contained the grasp type “Lateral,” there would be a 1 in row 1. The model works to minimize the sum of standard error for each β coefficient. Once optimal β coefficients have been found, an F-test is performed, followed by t-tests to determine which β coefficients are significantly influencing the model. We then perform a multiple comparison correction (False Detection Rate (FDR)) to correct p-values. A unit is tuned to the variable of interest if the corrected p-value is < 0.05.

This method was used to identify single-units tuned to each variable (grasp, size, shape) for each session. We could then determine which phases (Planning, Action, both), contexts (Hand, Hand+Object, Object, overlapping), or selectivity (selective (tuned to only one class within a variable) or mixed-selective (tuned to two or more classes)) these tuned units demonstrated.

### Classification and significance testing

A classifier was used to determine how well each feature could be differentiated within each phase, and throughout the entirety of the trial, using the neural firing rates recorded during each task. For each task, each session and each array were analyzed individually using linear discriminate analysis (LDA), assuming an identical diagonal covariance matrix for each set of variables. These assumptions, compared to a full diagonal covariance matrix, resulted in the best classification accuracies as determined by Wandelt et al. (2022)^56^. Classifiers were trained in two ways: 1) using z-scored average firing rate data from each phase (Cue: 2 seconds, Action: 4 seconds); 2) using z-scored average firing rate data from each 50 ms timebin throughout the entire trial. The 10 highest principal components (PCs), or the PCs explaining >90% of the variance (whichever was higher), were selected via principal component analysis (PCA) and served as feature selection on the training set. When fewer than 10 PCs were available, all features were used. Decoding performance was estimated using leave-one-out cross validation, and a 95% confidence interval was computed using the standard error of the mean.

To determine the significance of classification performance, we produced a null dataset by repeating classification with shuffled labels. 95% confidence intervals were computed for the null dataset and compared to the actual data. A t-test was used to compare the classification performances between actual and shuffled data. A false-detection rate correction was then applied and bins with p > 0.05 are determined. Bins were ultimately determined to be significant if there were ≥3 consecutive bins with p > 0.05 and denoted with a line above the bins.

### Planning and execution latencies and significance testing

Onset of Planning and Execution latencies were determined using the same classification method, with real and null datasets created by repeating classification with actual and shuffled labels 1000 times, respectively. Onset of Planning activity was determined by the first timebin to achieve significant grasp decoding compared to the null dataset. Significance was determined by performing a Mann-Whitney test between real and null classification accuracies at each timebin. Onset of execution was determined by training an LDA classifier to identify Go vs. No-Go trials, ensuring equal number of trials. The first timebin to achieve significant decoding compared to the null dataset was determined for each session and served as the execution latency of each region, determined by Mann-Whitney tests. Each analysis resulted in one datapoint for each region for every session. Datapoints that were deemed impossible (i.e., significance in the first timebin of Planning) were discarded. Two-sided Wilcoxon signed rank tests with FDR correction were performed to compare latencies among each region within subjects.

### Cross-phase and Cross-context classification and significance testing

To assess the shared neural activity between different task phases (cross-phase), we averaged neural activity across each phase and used the LDA classifier described above to train the classifier on one phase (i.e., Planning) and to test it on all phases (ITI, Planning, Action). This was repeated for each session individually, and a 95% confidence interval of the mean was calculated across sessions. A paired two-tailed t-test along with FDR correction of the p-vales was performed to determine significant differences in classification accuracies between phases.

P-values ≤0.05 are denoted with a * symbol, ≤0.01 with **, and ≤0.001 with ***. As we were predominantly interested in generalization between Planning and Action, those results are shown, along with ITI which serves as a control. The decoder was trained to distinguish grasp types during planning and tested on activity during Action (and vice versa). Generalization between phases occurs when training on one phase and testing on another results in classification accuracies above chance. Generalization can be complete or incomplete. Complete generalization occurs when there is no significant difference in the performance of the classifier when tested on the matched phase (i.e., train: Planning, test: Planning) or unmatched (i.e., train: Planning, test: Action), suggesting a similar representation of grasp in each phase. Incomplete generalization occurs when there is a significant difference between phases, suggesting an altered representation of grasp.

The same method was used to assess the shared neural activity across different contexts (cross-context), this time training the classifier on one context (i.e., Hand only) and testing it on all contexts (Hand only, Hand+Object, Object only). Significance was determined and denoted as above. From these statistics, we determined whether contexts generalized to one another, either completely (no significant difference (+)) or incompletely (significantly different but above chance (-)).

Due to the computational requirements of these analyses (cross-validation partitioning within each phase and context) and necessary limiting of trials per session for participant comfort, these analyses could not be successfully implemented for participant S2.

### Neuron Dropping Curves

Neuron dropping curves (NDCs) were computed using a neuron grid containing the first 5 units, the 7^th^, then increments of 3 units until reaching the unit minimum number of units for each region. The lowest number of units recorded in a single session was identified to ensure that all sessions were included in the analysis. The average firing rate for each channel was calculated and z-scored for each phase. For each step, we randomly selected unit(s) then partitioned with a K-fold of 8. We trained an LDA classifier with cross validation and averaged across folds. We repeated this 50 times for each neuron count, obtaining the expected decoding accuracy using k randomly sampled neurons. NDCs were calculated for each session and then averaged across sessions. Peak classification accuracy in **Fig. 2D,F,G** were determined using the peak accuracies from the NDCs.

---

**Supplementary Figure 1.**
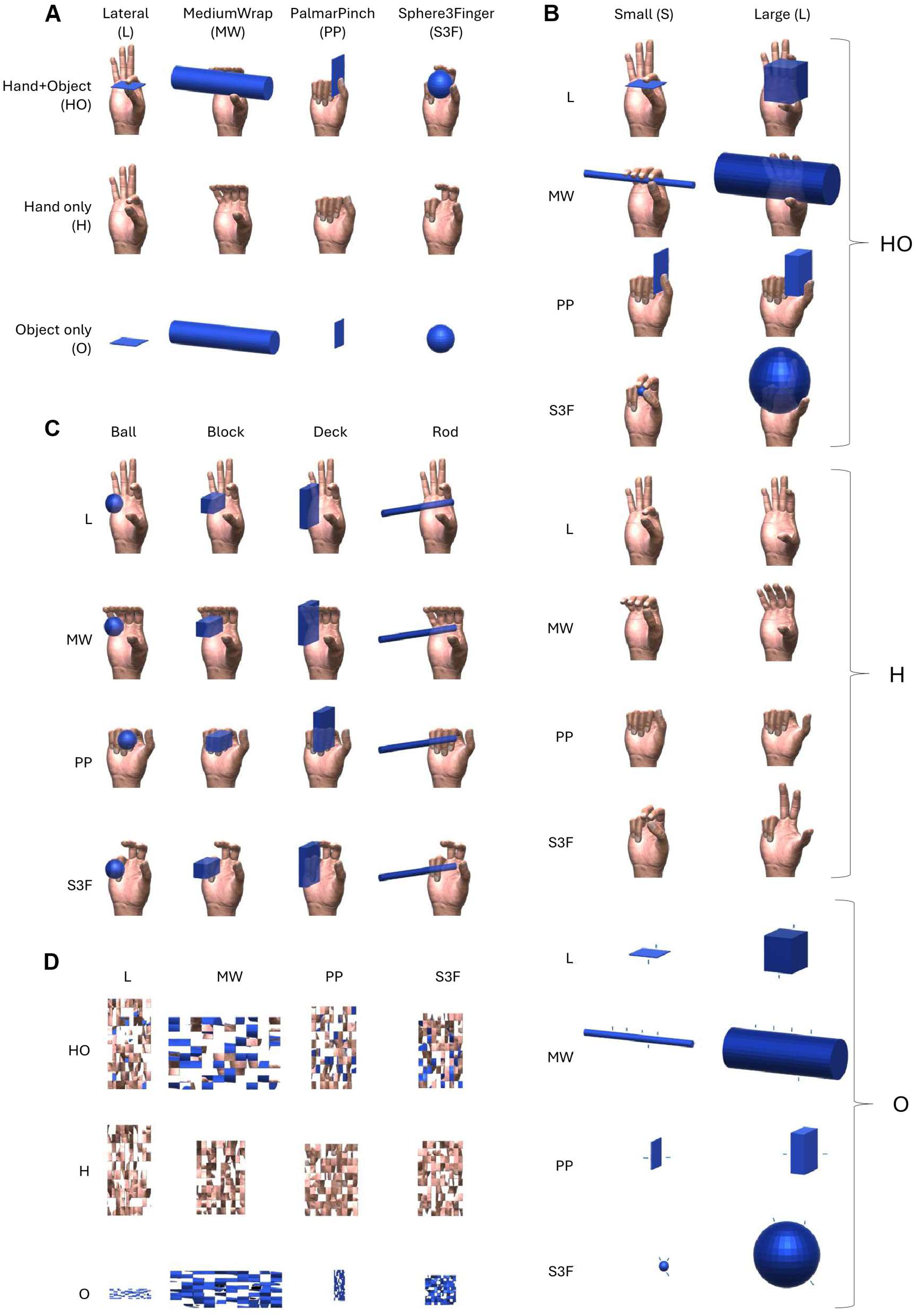
All variations of cued static images. A) All grasp/context combinations. B) All grasp/size/context combinations for Varied Sizes Task. Blue dashes added to the Object only context to indicate to participants which fingers to use and where they should imagine grasping. C) Static images for Disassociation Task depicting each grasp paired with each object (object dimensions were altered to enable feasible grasping by each grasp type). D) Pixelated images used as visual control.

**Supplementary Figure 2.**
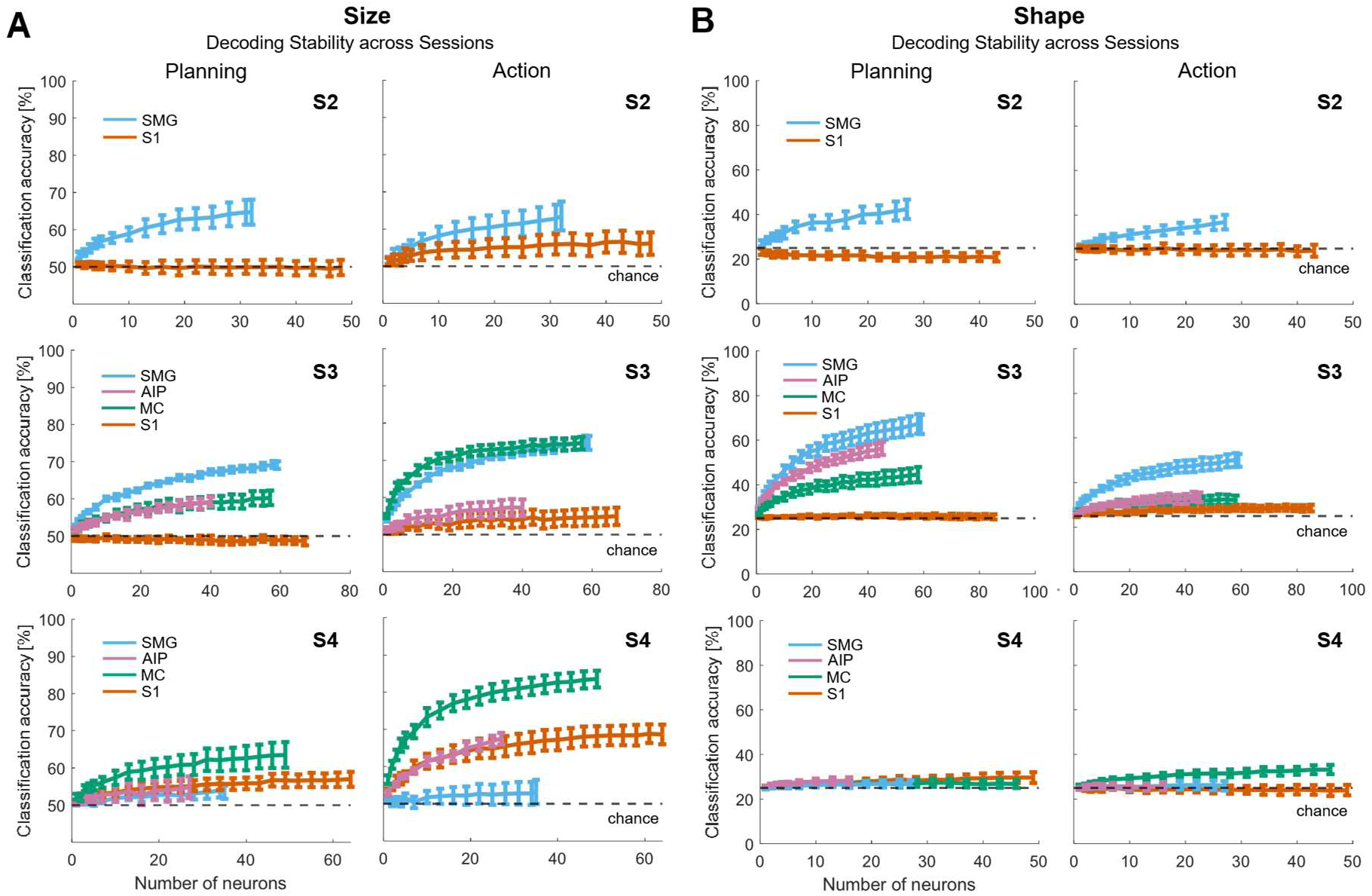
Neuron Dropping Curves for classifying object size (A) and shape (B) in each participant during Planning and Action phases

**Supplementary Fig. 3:**
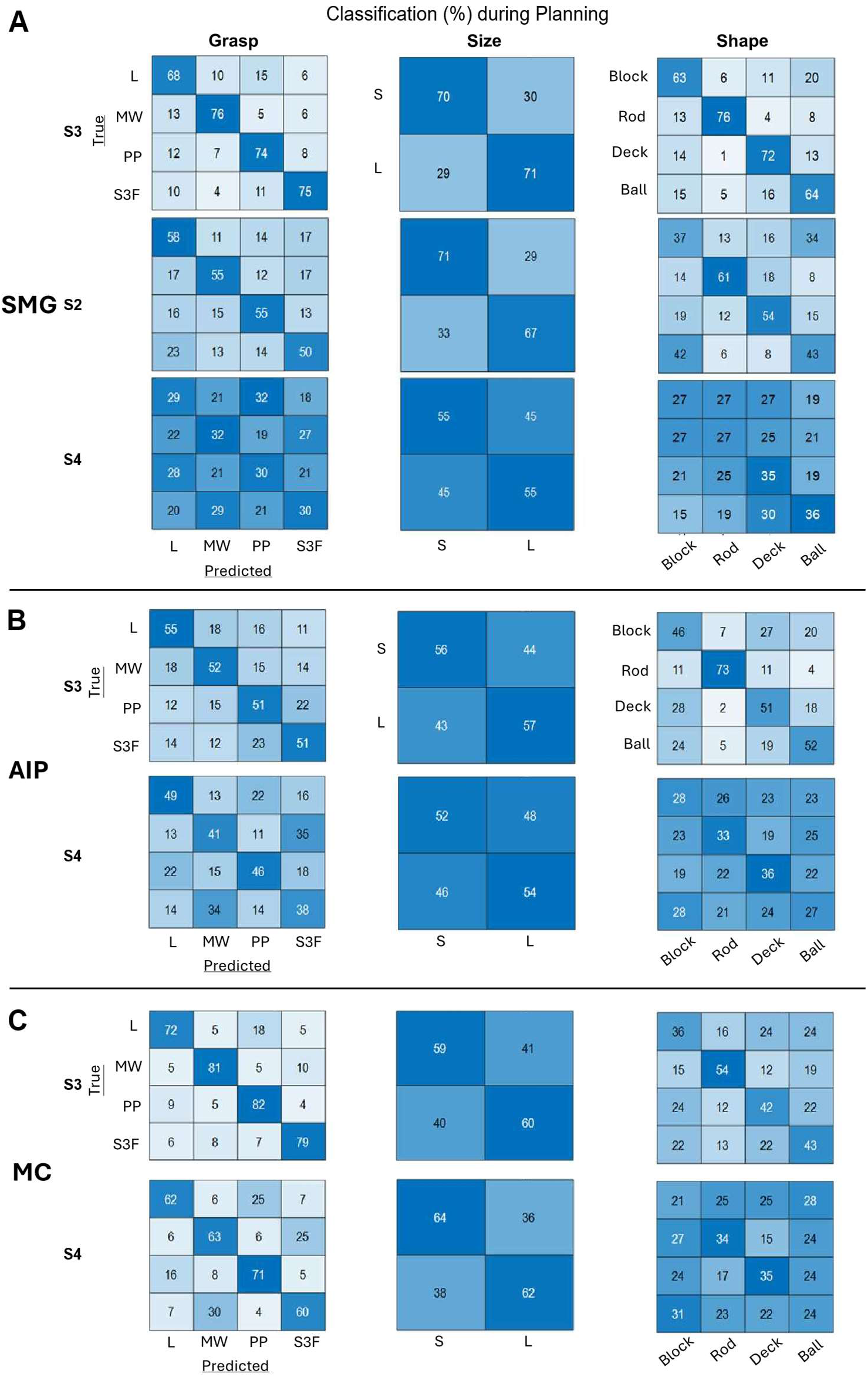
Classification accuracies (%) for each grasp type (column 1), size (col. 2), and shape (col. 3) from SMG (A), AIP (B), and MC (C) during Planning phase in all participants.

**Supplementary Fig. 4:**
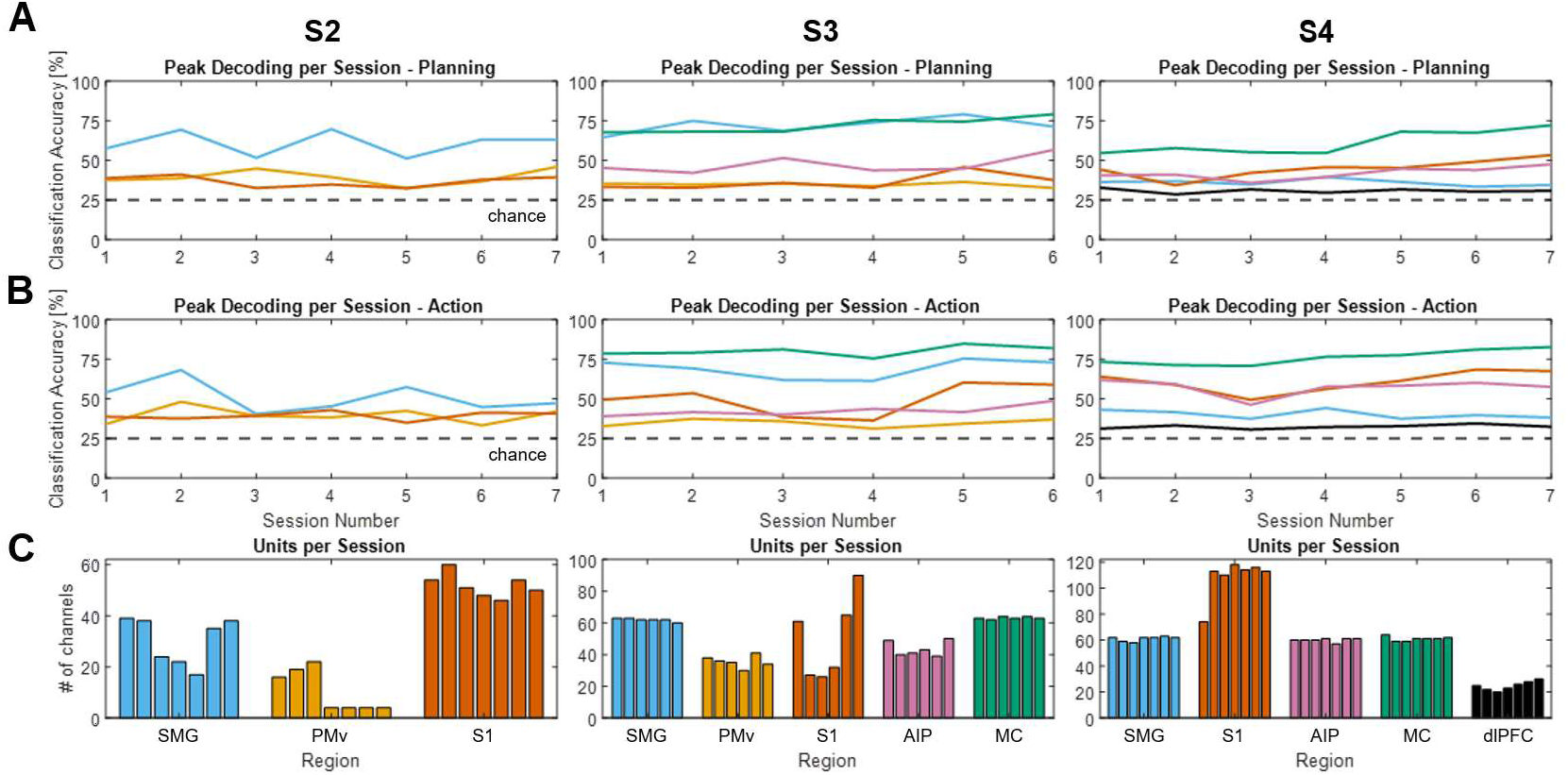
Decoding stability across sessions. Peak grasp decoding performance for each region during each session determined by the highest classification accuracy in a single timebin during Planning (A) and Action (B) phases. Each line is a region, colors correspond to C. C) Number of channels across sessions is relatively stable and consistent within and between participants. Channels are channels that exhibited multi-unit activity during the task as determined by pre-processing threshold crossings (see Methods).

**Supplementary Fig. 5:**
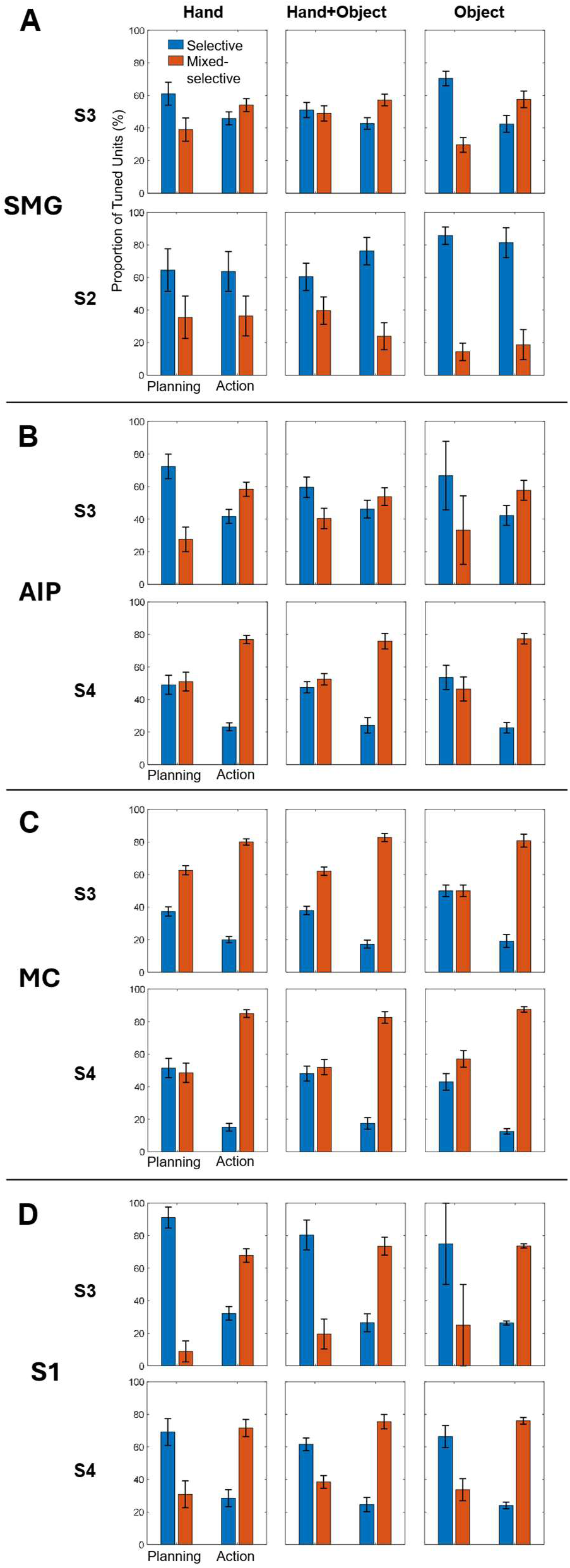
Selective vs. Mixed-selective grasp-tuned units during Planning and Action in SMG (A), AIP (B), MC (C), and S1 (D) during each context: Hand only (col. 1), Hand+Object (col. 2), and Object only (col. 3).

**Supplementary Fig. 6:**
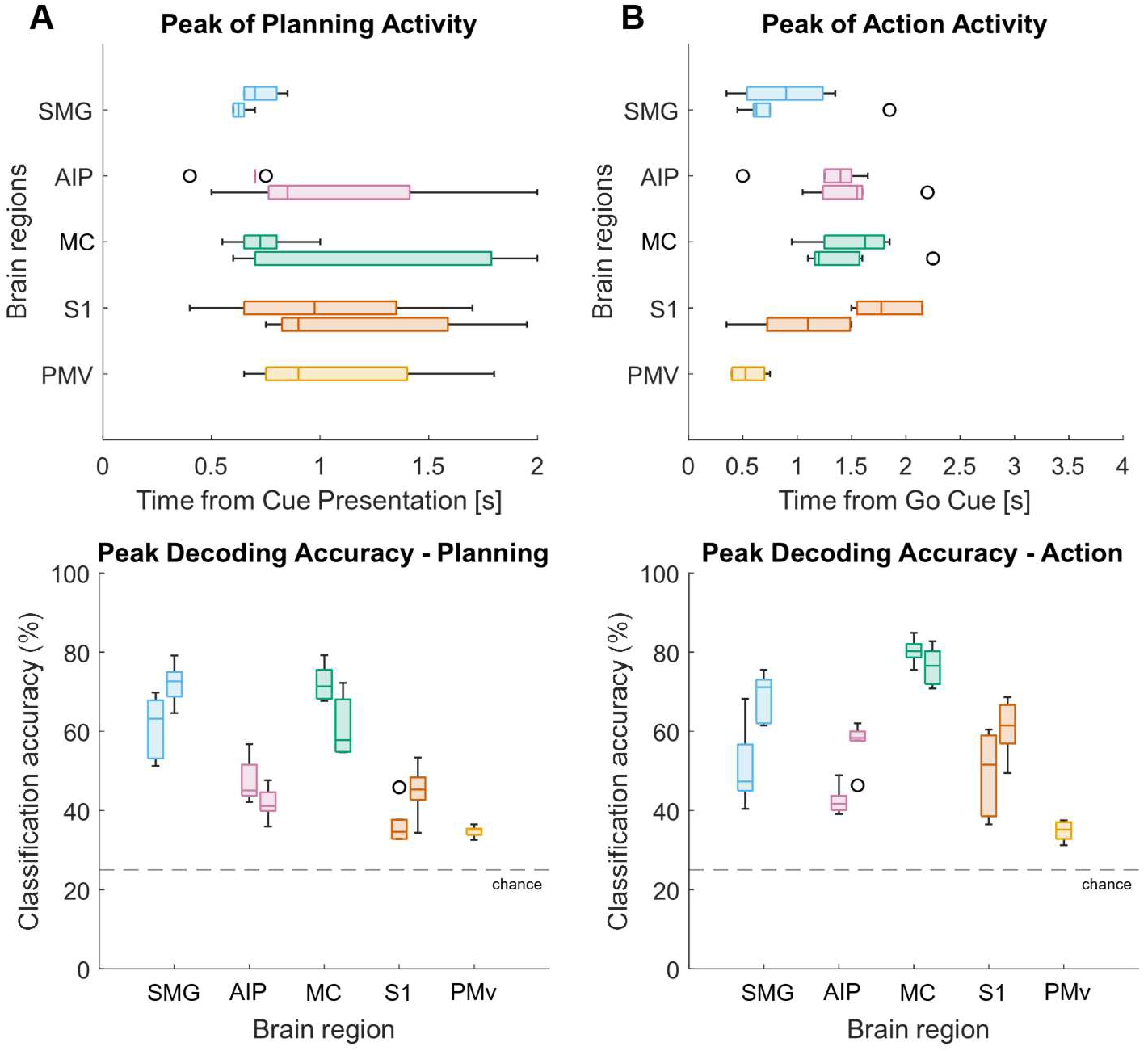
Peak classification timing and accuracies across regions and participants. A) During Planning: SMG peaks at 681 ms (S2,S3) with 67% average, AIP peaks at 854.1667 ms (S3,S4) with 45% average, M1 peaks at 945.8333 ms (S3,S4) with 67% average, S1 peaks at 1094 ms (S3,S4) with 41% average, PMv peaks at 1066.7 ms (S3) with 35% average. B) During Action: SMG peaks at 851 ms (S2,S3) with 60% average, AIP peaks at 1402 ms (S3,S4) with 50% average, M1 peaks at 1473 ms (S3,S4) with 78% average, S1 peaks at 1437 ms (S3,S4) with 55% average, PMv peaks at 550 ms (S3) with 35% average. Each boxplot is one participant.

**Supplementary Figure 7.**
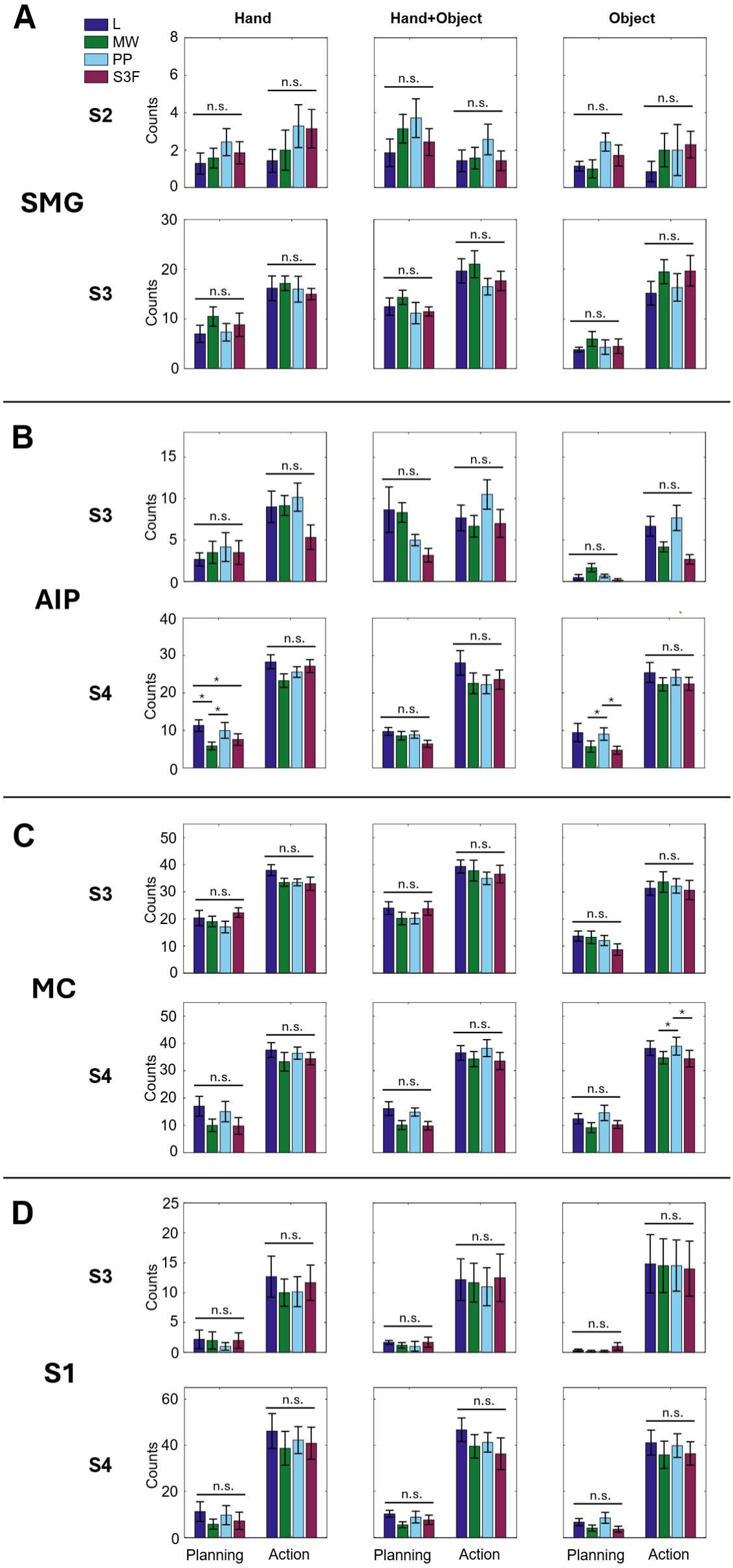
Number of units tuned to specific grasp types averaged across sessions for each participant in SMG (A), AIP (B), MC (C), and S1 (D). P-values ≤0.05 are denoted with a * symbol, ≤0.01 with **, and ≤0.001 with ***

**Supplementary Figure 8.**
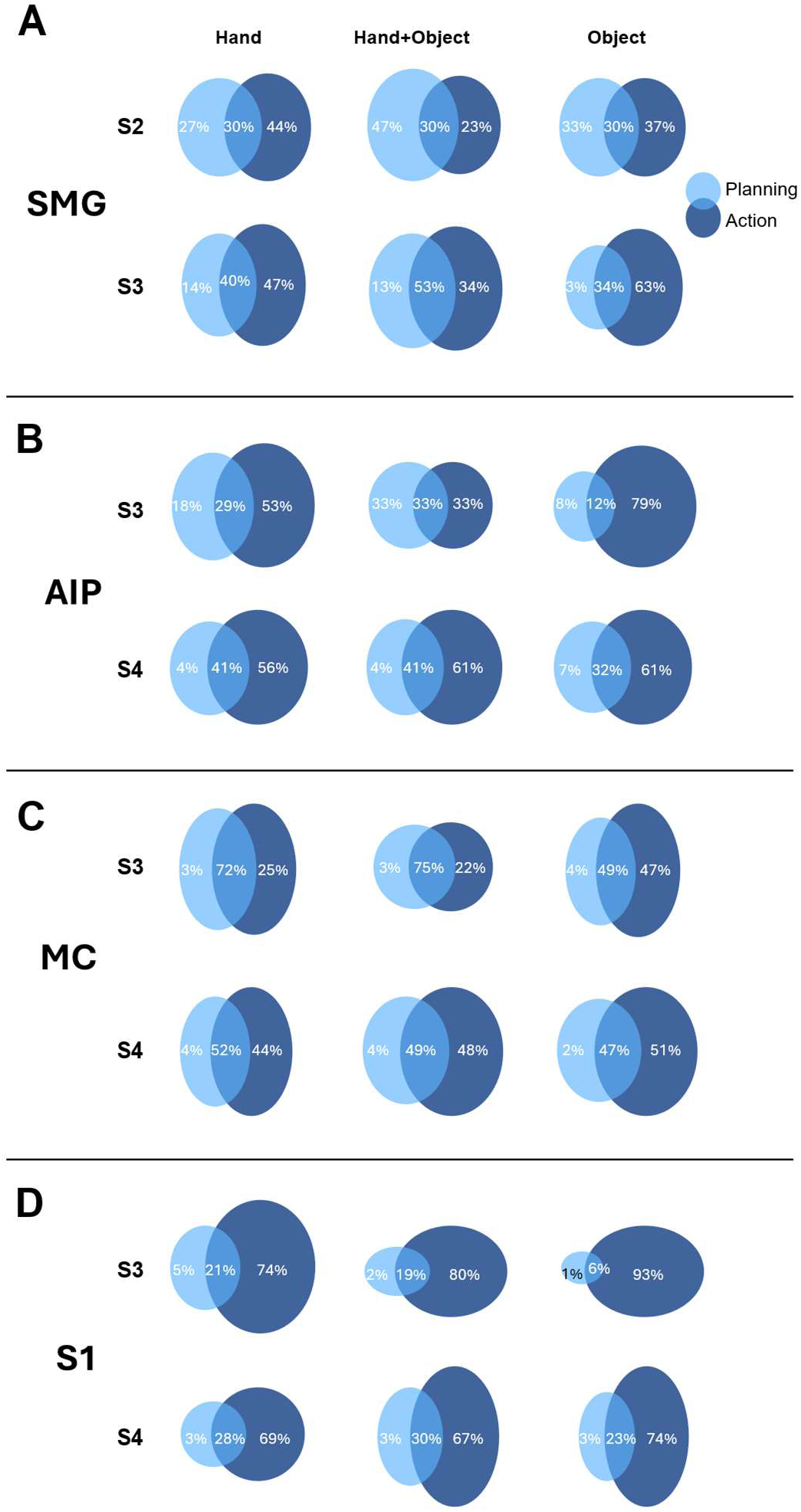
Proportion of grasp-tuned units during Planning (light blue) and Action (dark blue) for each participant in SMG (A), AIP (B), MC (C), and S1 (D).

**Supplementary Fig. 9:**
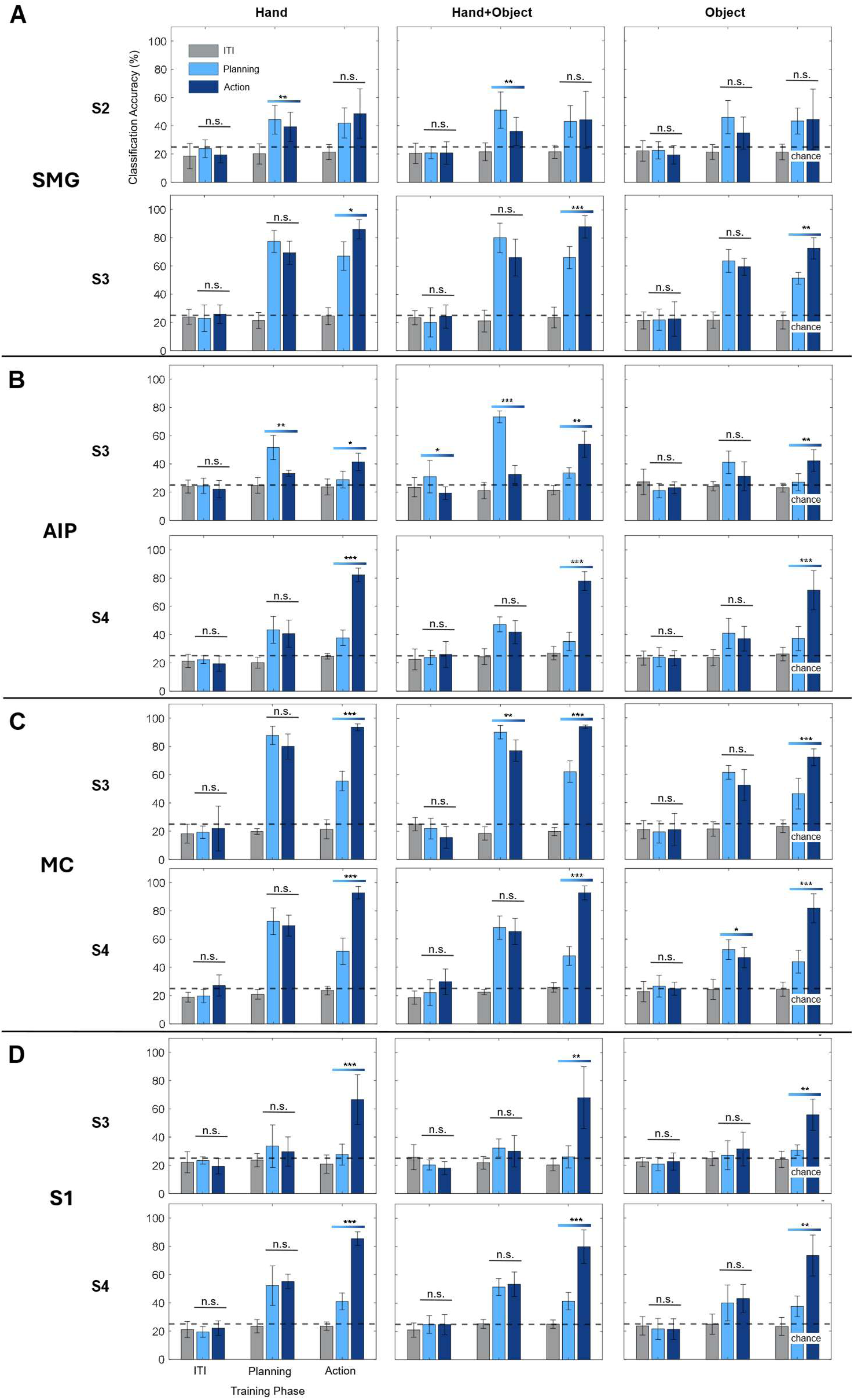
Cross-phase classification separated by each context (Hand only (col. 1), Hand+Object (col. 2), and Object only (col. 3)) in SMG (A), AIP (B), MC (C), and S1 (D). Planning does generalize to Action, but Action does not generalize to Planning. P-values ≤0.05 are denoted with a * symbol, ≤0.01 with **, and ≤0.001 with ***.

**Supplementary Fig. 10:**
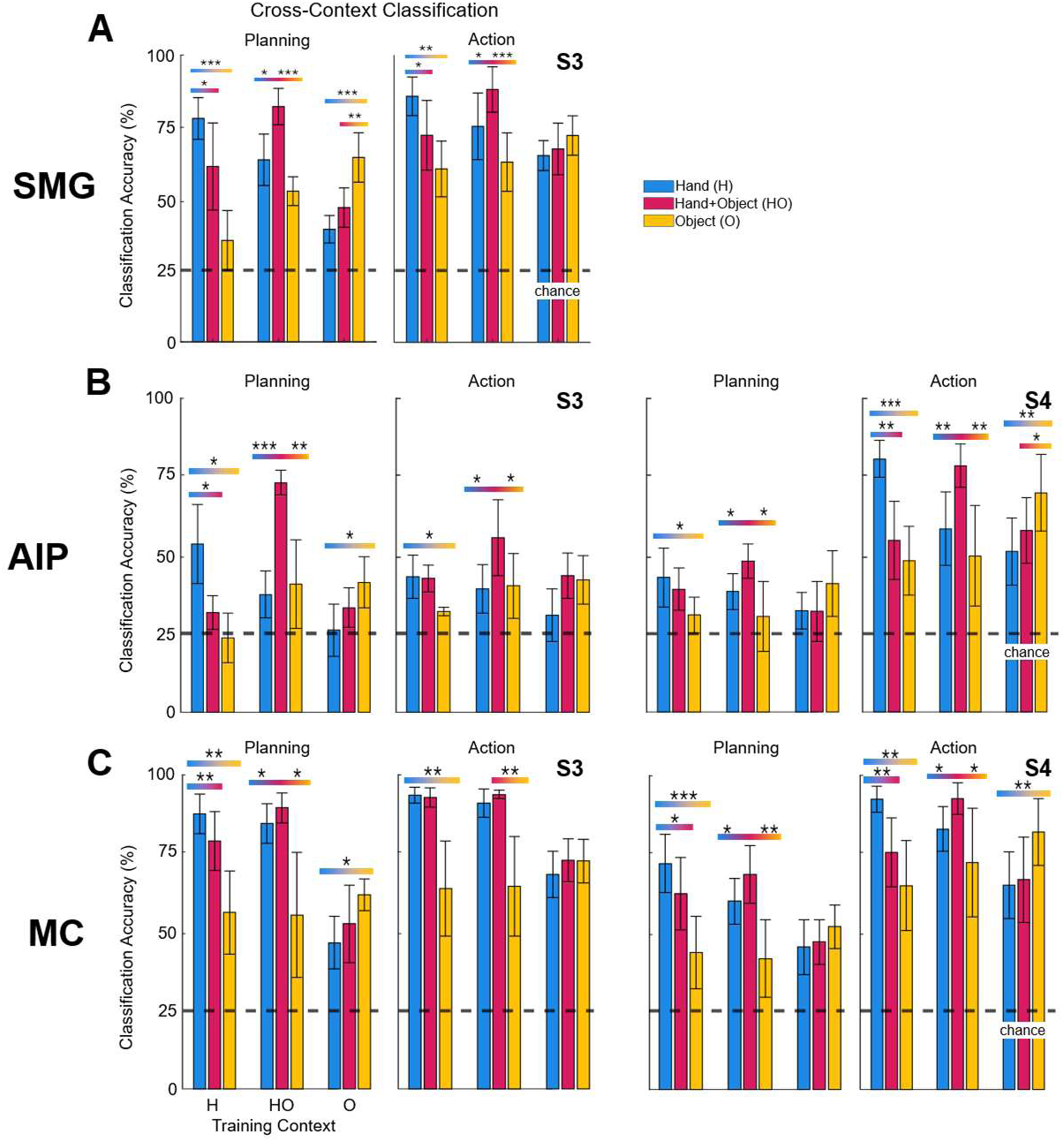
Cross-context classification during Planning and Action phases for each participant (S2 excluded, see Methods) in SMG (A), AIP (B), and MC (C). P-values ≤0.05 are denoted with a * symbol, ≤0.01 with **, and ≤0.001 with ***.

**Supplementary Fig. 11:**
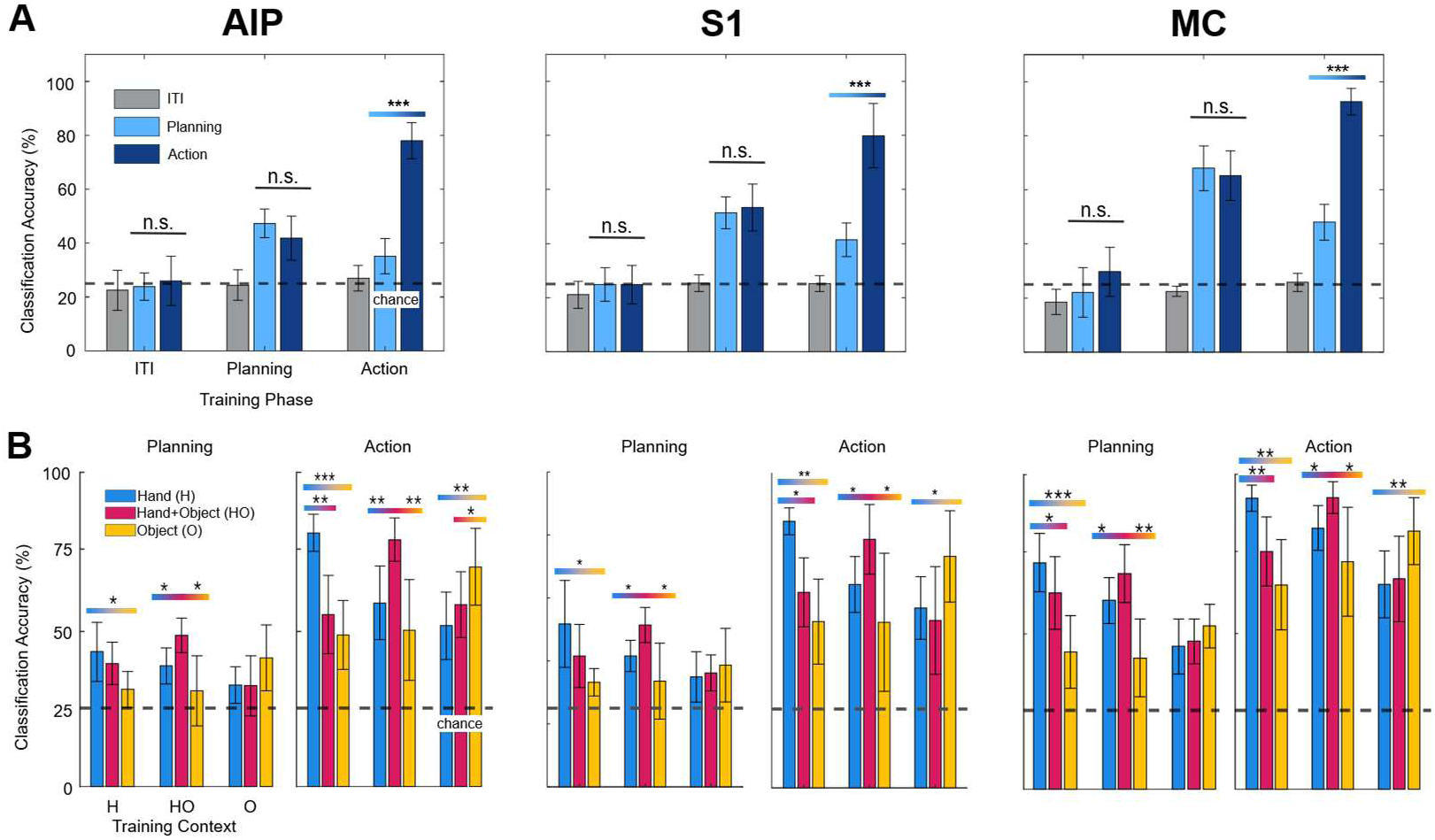
S4’s AIP has similar response to primary sensorimotor areas. A) Cross-phase classification reveals a similar pattern of complete generalization from Planning to Action, but incomplete generalization from Action to Planning in AIP (left), S1 (middle), and MC (right). Example of HO shown. B) Cross-context classification reveals markedly similar classification patterns during Planning and Action, with each context reaching similar classification accuracies. During Planning, training on HO consistently leads to incomplete generalization. During Action, we observe predominantly incomplete generalization of each context.

**Supplementary Fig. 12:**
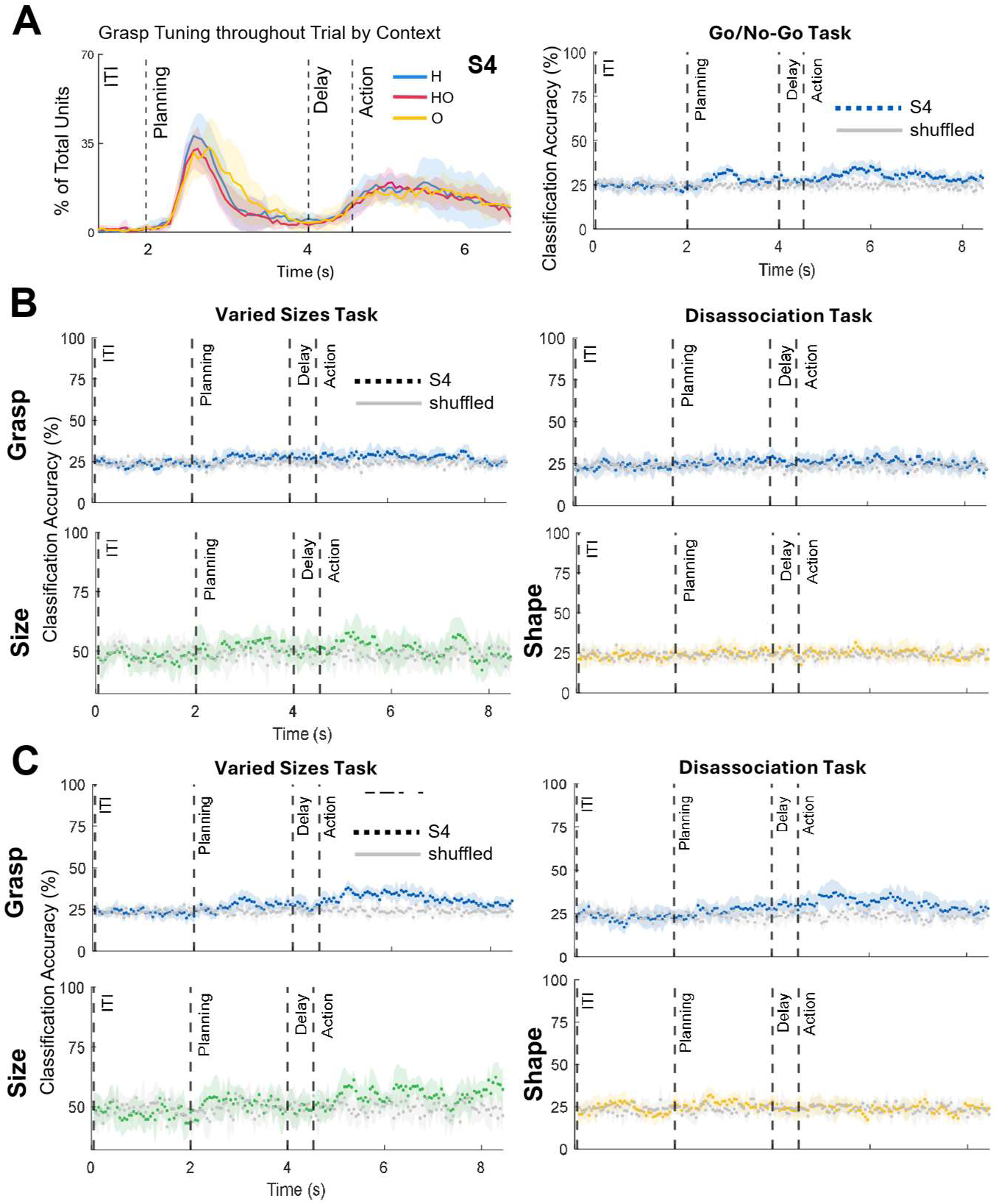
S4’s tuning and classification results in SMG and AIP. A) S4’s SMG array demonstrates tuning to grasp in each context (right), but no significant grasp classification (left). B) Classification of grasp (top), size (bottom right), and shape (bottom left) from S4’s SMG array are all chance level. These results are consistent for every task. C) Classification of grasp (top), size (bottom right), and shape (bottom left) from S4’s AIP array are nearly all chance level. Some residual grasp decoding appears in the Varied Sizes task, but this is drastically decreased from the Go/No-Go task accuracies and is completely non-significant during the Disassociation task.

**Supplementary Fig. 13:**
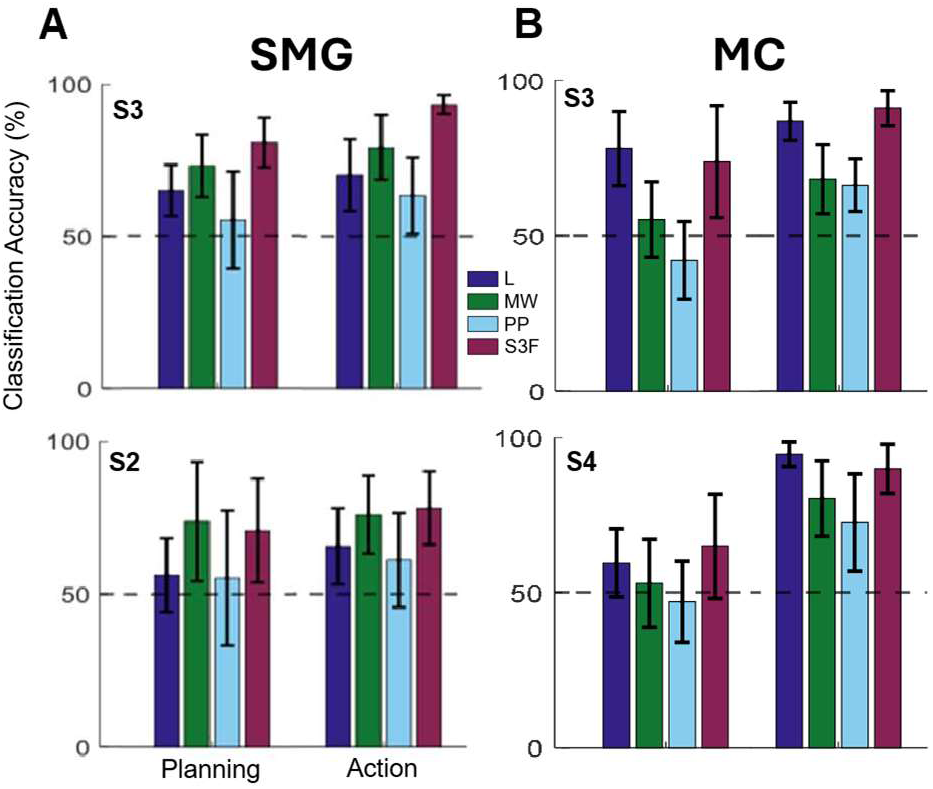
Size classification accuracies per grasp type in SMG (A) and MC (B) during Planning and Action demonstrate higher classification accuracies of size within specific grasp types.

**Supplementary Figure 14.**
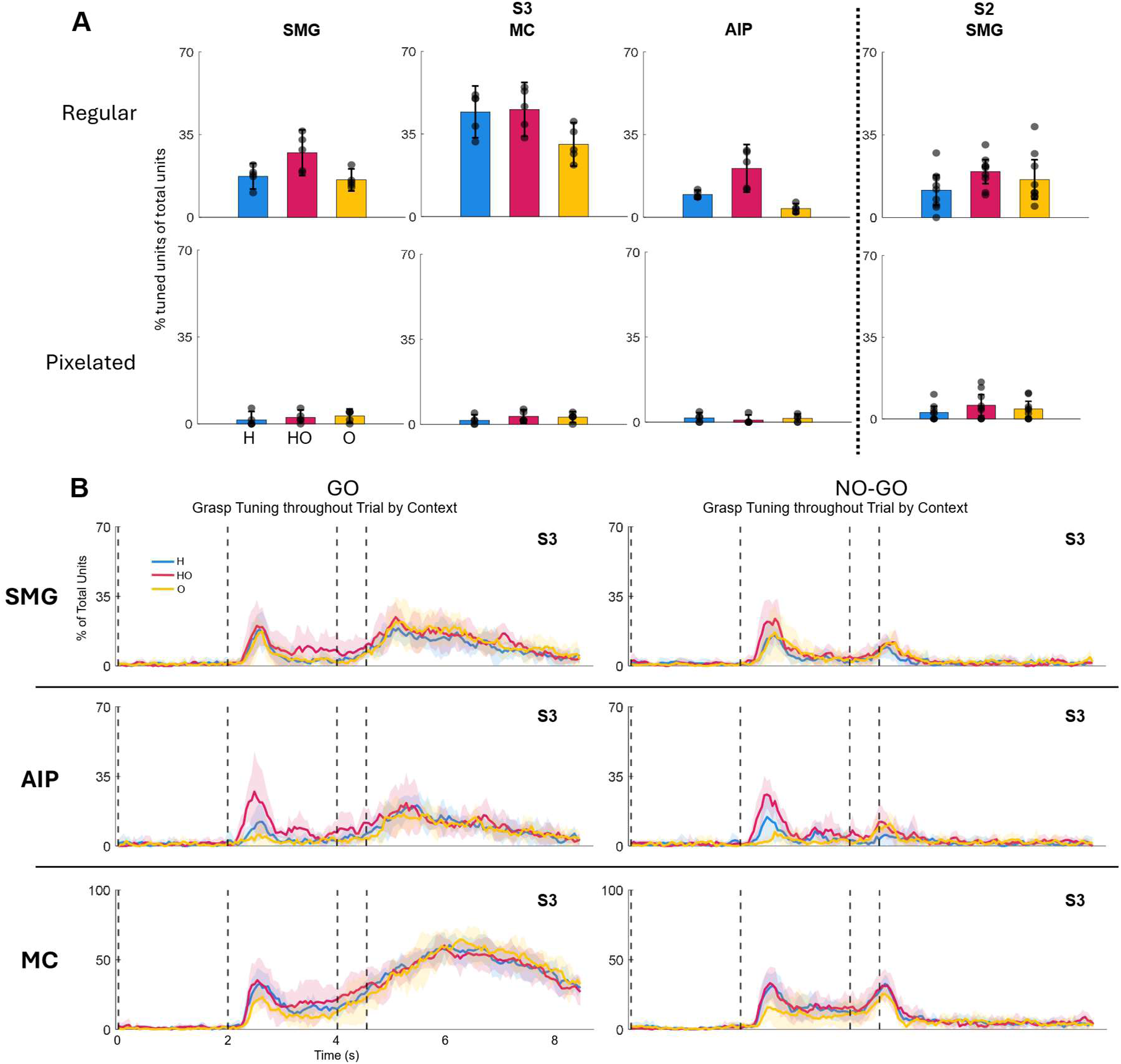
Control experiments demonstrate that participants are performing the task appropriately and the results are due to motor planning, not visual stimuli. A) Tuning to pixelated images during the Planning phase show that very few units within each region are tuned to purely visual stimuli. B) No-Go trials result in similar tuning level until the Action phase, where tuned units precipitously drop off. Data shown here are from participant S3, but the result is the same among all participants.

## REFERENCES

1. Edemekong, P. F., Bomgaars, D. L., Sukumaran, S. & Schoo, C. Activities of Daily Living. in StatPearls (StatPearls Publishing, Treasure Island (FL), 2025).

2. Anderson, K. D. Targeting recovery: priorities of the spinal cord-injured population. J Neurotrauma 21, 1371–1383 (2004).

3. Collinger, J. L. et al. 7 degree-of-freedom neuroprosthetic control by an individual with tetraplegia. Lancet 381, 557–564 (2013).

4. Nicolas-Alonso, L. F. & Gomez-Gil, J. Brain Computer Interfaces, a Review. Sensors 12, 1211–1279 (2012).

5. Schaffelhofer, S. & Scherberger, H. Object vision to hand action in macaque parietal, premotor, and motor cortices. eLife 5, (2016).

6. Michaels, J. A., Schaffelhofer, S., Agudelo-Toro, A. & Scherberger, H. A goal-driven modular neural network predicts parietofrontal neural dynamics during grasping. PNAS 117, 32124–32135 (2020).

7. Schaffelhofer, S., Agudelo-Toro, A. & Scherberger, H. Decoding a Wide Range of Hand Configurations from Macaque Motor, Premotor, and Parietal Cortices. J. Neurosci. 35, 1068– 1081 (2015).

8. Davare, M., Kraskov, A., Rothwell, J. C. & Lemon, R. N. Interactions between areas of the cortical grasping network. Current Opinion in Neurobiology 21, 565–570 (2011).

9. Muir, R. B. & Lemon, R. N. Corticospinal neurons with a special role in precision grip. Brain Research 261, 312–316 (1983).

10. Georgopoulos, A. P., Kalaska, J. F., Caminiti, R. & Massey, J. T. On the relations between the direction of two-dimensional arm movements and cell discharge in primate motor cortex. J Neurosci 2, 1527–1537 (1982).

11. Yan, Y. et al. Evolution of object identity information in sensorimotor cortex throughout grasp. Nat Commun 17, 2784 (2026).

12. Snyder, L. H., Batista, A. P. & Andersen, R. A. Intention-related activity in the posterior parietal cortex: a review. Vision Res 40, 1433–1441 (2000).

13. Townsend, B. R., Subasi, E. & Scherberger, H. Grasp Movement Decoding from Premotor and Parietal Cortex. J. Neurosci. 31, 14386–14398 (2011).

14. Yokoi, A., Arbuckle, S. A. & Diedrichsen, J. The Role of Human Primary Motor Cortex in the Production of Skilled Finger Sequences. J Neurosci 38, 1430–1442 (2018).

15. Cheney, P. D. Role of cerebral cortex in voluntary movements. A review. Phys Ther 65, 624– 635 (1985).

16. Vingerhoets, G. Contribution of the posterior parietal cortex in reaching, grasping, and using objects and tools. Front. Psychol. 5, (2014).

17. Osiurak, F. & Badets, A. Tool use and affordance: Manipulation-based versus reasoning- based approaches. Psychological Review 123, 534–568 (2016).

18. Orban, G. A. & Caruana, F. The neural basis of human tool use. Front Psychol 5, 310 (2014).

19. McDowell, T., Holmes, N. P., Sunderland, A. & Schürmann, M. TMS over the supramarginal gyrus delays selection of appropriate grasp orientation during reaching and grasping tools for use. Cortex 103, 117–129 (2018).

20. Wandelt, S. K. et al. Decoding grasp and speech signals from the cortical grasp circuit in a tetraplegic human. Neuron 110, 1777–1787.e3 (2022).

21. Buchwald, M., Przybylski, Ł. & Króliczak, G. Decoding Brain States for Planning Functional Grasps of Tools: A Functional Magnetic Resonance Imaging Multivoxel Pattern Analysis Study. Journal of the International Neuropsychological Society 24, 1013–1025 (2018).

22. Filimon, F., Nelson, J. D., Huang, R.-S. & Sereno, M. I. Multiple Parietal Reach Regions in Humans: Cortical Representations for Visual and Proprioceptive Feedback during On-Line Reaching. J Neurosci 29, 2961–2971 (2009).

23. Wandelt, S. K. et al. Representation of internal speech by single neurons in human supramarginal gyrus. Nat Hum Behav 8, 1136–1149 (2024).

24. Zhao, Y., Hessburg, J. P., Asok Kumar, J. N. & Francis, J. T. Paradigm Shift in Sensorimotor Control Research and Brain Machine Interface Control: The Influence of Context on Sensorimotor Representations. Front. Neurosci. 12, (2018).

25. Hochberg, L. R. et al. Reach and grasp by people with tetraplegia using a neurally controlled robotic arm. Nature 485, 372–375 (2012).

26. Flesher, S. N. et al. A brain-computer interface that evokes tactile sensations improves robotic arm control. Science 372, 831–836 (2021).

27. Downey, J. E. et al. Motor cortical activity changes during neuroprosthetic-controlled object interaction. Sci Rep 7, 16947 (2017).

28. Feix, T., Romero, J., Schmiedmayer, H.-B., Dollar, A. M. & Kragic, D. The GRASP Taxonomy of Human Grasp Types. IEEE Trans. Human-Mach. Syst. 46, 66–77 (2016).

29. Aflalo, T. et al. Neural prosthetic context profoundly shapes single-neuron responses while preserving the functional distinctions between human motor and posterior parietal cortices. https://www.abstractsonline.com/pp8/#!/10485/presentation/10264 (2021).

30. Bjånes, D. A. et al. Charge density of multi-channel intra-cortical micro-stimulation modulates intensity and naturalness of evoked somatosensations. J Neural Eng 22, (2025).

31. Jafari, M. et al. The human primary somatosensory cortex encodes imagined movement in the absence of sensory information. Commun Biol 3, 757 (2020).

32. Bjånes, D. A. et al. Quantifying physical degradation alongside recording and stimulation performance of 980 intracortical microelectrodes chronically implanted in three humans for 956-2130 days. Acta Biomaterialia 198, 188–206 (2025).

33. Yan, Y. et al. Evolution of object identity information in sensorimotor cortex throughout grasp. Nat Commun 10.1038/s41467-026-69502-0 (2026) doi:10.1038/s41467-026-69502-0.

34. Gallivan, J. P., McLean, D. A., Valyear, K. F., Pettypiece, C. E. & Culham, J. C. Decoding Action Intentions from Preparatory Brain Activity in Human Parieto-Frontal Networks. J. Neurosci. 31, 9599–9610 (2011).

35. Ariani, G., Shahbazi, M. & Diedrichsen, J. Cortical Areas for Planning Sequences before and during Movement. J. Neurosci. 45, (2025).

36. Andersen, R. A., Aflalo, T., Bashford, L., Bjånes, D. & Kellis, S. Exploring Cognition with Brain–Machine Interfaces. Annual Review of Psychology 73, 131–158 (2022).

37. Tien, R. N. Encoding of Object Presence and Manipulation Affordances in the Frontoparietal Grasp Network. https://d-scholarship.pitt.edu/40183 (2021).

38. Wodlinger, B. et al. Ten-dimensional anthropomorphic arm control in a human brain−machine interface: difficulties, solutions, and limitations. J. Neural Eng. 12, 016011 (2014).

39. Tortolani, A. F. et al. How different immersive environments affect intracortical brain computer interfaces. J Neural Eng 22, 016032 (2025).

40. Guthrie, M. D. et al. The impact of distractions on intracortical brain–computer interface control of a robotic arm. Brain-Computer Interfaces 9, 23–35 (2022).

41. Velliste, M. et al. Motor Cortical Correlates of Arm Resting in the Context of a Reaching Task and Implications for Prosthetic Control. J. Neurosci. 34, 6011–6022 (2014).

42. Mender, M. J. et al. The impact of task context on predicting finger movements in a brain- machine interface. eLife 12, e82598 (2023).

43. Stavisky, S. D. et al. Neural ensemble dynamics in dorsal motor cortex during speech in people with paralysis. eLife 8, e46015 (2019).

44. Shaikhouni, A., Donoghue, J. P. & Hochberg, L. R. Somatosensory responses in a human motor cortex. J Neurophysiol 109, 2192–2204 (2013).

45. Shelchkova, N. D. et al. Microstimulation of human somatosensory cortex evokes task- dependent, spatially patterned responses in motor cortex. Nat Commun 14, 7270 (2023).

46. Vargas-Irwin, C. E. et al. Watch, Imagine, Attempt: Motor Cortex Single-Unit Activity Reveals Context-Dependent Movement Encoding in Humans With Tetraplegia. Front. Hum. Neurosci. 12, (2018).

47. Kunz, E. M. et al. Inner speech in motor cortex and implications for speech neuroprostheses. Cell 188, 4658–4673.e17 (2025).

48. Aflalo, T. et al. A shared neural substrate for action verbs and observed actions in human posterior parietal cortex. Sci Adv 6, eabb3984 (2020).

49. Bougou, V. et al. Hierarchical and Context-Dependent Encoding of Actions in Human Posterior Parietal and Motor Cortex. Preprint at 10.1101/2025.11.10.687245 (2025).

50. Kunigk, N. G. et al. Motor somatotopy impacts imagery strategy success in human intracortical brain–computer interfaces. J. Neural Eng. 22, 026004 (2025).

51. Aflalo, T. et al. Decoding motor imagery from the posterior parietal cortex of a tetraplegic human. Science 348, 906–910 (2015).

52. Messier, J. & Kalaska, J. F. Covariation of primate dorsal premotor cell activity with direction and amplitude during a memorized-delay reaching task. J Neurophysiol 84, 152– 165 (2000).

53. Baud-Bovy, G. & Soechting, J. F. Two virtual fingers in the control of the tripod grasp. J Neurophysiol 86, 604–615 (2001).

54. Santello, M., Baud-Bovy, G. & Jörntell, H. Neural bases of hand synergies. Front Comput Neurosci 7, 23 (2013).

55. Murata, A., Gallese, V., Luppino, G., Kaseda, M. & Sakata, H. Selectivity for the shape, size, and orientation of objects for grasping in neurons of monkey parietal area AIP. J Neurophysiol 83, 2580–2601 (2000).

56. Wandelt, S. K. et al. Online internal speech decoding from single neurons in a human participant. 2022.11.02.22281775 Preprint at 10.1101/2022.11.02.22281775 (2022).

57. Marzke, M. W., Wullstein, K. L. & Viegas, S. F. Evolution of the power (“squeeze”) grip and its morphological correlates in hominids. 10.1002/ajpa.1330890303 doi:10.1002/ajpa.1330890303.

58. Bardo, A. et al. The Precision of the Human Hand: Variability in Pinch Strength and Manual Dexterity. Symmetry 14, 71 (2022).

59. Rilling, J. K. Comparative primate neuroimaging: insights into human brain evolution. Trends in Cognitive Sciences 18, 46–55 (2014).

60. Wey, H.-Y. et al. Multi-region hemispheric specialization differentiates human from nonhuman primate brain function. Brain Struct Funct 219, 2187–2194 (2014).

61. Pang, J. C., Rilling, J. K., Roberts, J. A., Heuvel, M. P. van den & Cocchi, L. Evolutionary shaping of human brain dynamics. eLife 11, e80627 (2022).

62. Mynhier, N. A. et al. The Compositional Encoding of Hand-Eye Coordinated Movements for Single Neurons in the Posterior Parietal Cortex. 2026.04.04.716384 Preprint at 10.64898/2026.04.04.716384 (2026).

63. Rokni, U., Steinberg, O., Vaadia, E. & Sompolinsky, H. Cortical representation of bimanual movements. J Neurosci 23, 11577–11586 (2003).

64. Downey, J. E. et al. The Motor Cortex Has Independent Representations for Ipsilateral and Contralateral Arm Movements But Correlated Representations for Grasping. Cereb Cortex 30, 5400–5409 (2020).

65. Oberhuber, M. et al. Four Functionally Distinct Regions in the Left Supramarginal Gyrus Support Word Processing. Cereb Cortex 26, 4212–4226 (2016).

66. Yamazaki, Y., Hashimoto, T. & Iriki, A. The posterior parietal cortex and non-spatial cognition. F1000 Biol Rep 1, 74 (2009).

